# Skill Memory Expansion Shapes Micro-Scale Dynamics of Skill Learning

**DOI:** 10.1101/2025.11.13.686780

**Authors:** Fumiaki Iwane, William Hayward, I. M. Dushyanthi Karunathilake, Ethan R. Buch, Leonardo G. Cohen

**Affiliations:** Human Cortical Physiology and Neurorehabilitation Section, National Institute of Neurological Disorders and Stroke, NIH, 9000 Rockville Pike, Bethesda, MD 20817, USA

**Author notes:** **Corresponding authors:** Leonardo G Cohen. **Lead Contact:** Leonardo G Cohen.

## Abstract

Motor skill learning depends on refining action sequences through practice, yet how learning unfolds moment-to-moment within trials remains unclear. Here, healthy participants learned generative sequential keypress skills at different speeds. Within trials, we identified reproducible high initial skill (HIS) segments, brief epochs of elevated performance followed by decline, that emerged only during learning of repeating sequences and were absent during practice of pseudo-randomized, non-repeating sequences. HIS keypress content scaled with execution speed and increased with practice. This expansion of HIS segment content continued after overall performance plateaued. Neural analyses showed increased theta–gamma phase-amplitude coupling (θ/γ PAC) and beta-band bursts during HIS segments. Hippocampal θ/γ PAC predicted HIS content, suggesting a mechanism for binding successive actions into progressively larger skill-memory representations during early learning. Together, these findings identify expansion of skill memory, rather than fatigue, as a key driver of early skill learning.

## Introduction

Motor learning, from typing to musical performance, relies on the execution of temporally precise action sequences, which constitute the core elements of fine motor expertise ^1 2 3–6^. During the early stages of acquiring challenging skills, performance improves rapidly, with gains arising predominantly across brief rest periods interleaved with practice (micro-offline intervals)^1,7–12^. Behavioral manipulations showed that retroactive interference immediately after practice reduces learning rates relative to interference imposed after a delay, consistent with a time-dependent stabilization of the motor memory ^13^. Micro-offline improvements are robust and reproducible and persist even when practice bouts are shortened to as little as 5 seconds in duration, ruling out simple recovery from performance fatigue as their primary source ^8,11,13–15^. Instead, converging evidence across species and techniques implicates rapid neural replay during the brief rest breaks in this offline memory consolidation process. In humans, the rate of hippocampo–neocortical replay measured with MEG ^7^ and the density of hippocampal sharp-wave ripples (80–120 Hz), a canonical marker of replay, measured with intracranial EEG ^14,15^ predict micro-offline improvements during early learning. Griffin et al. extended these correlational findings to rhesus macaques^16^. Importantly, the same study provided the first causal evidence for this relationship as 20-Hz AC stimulation effectively disrupted ripple-embedded replay activity in the primary motor cortex, eliminated micro-offline performance gains in subsequent rest breaks (analogous to those interspersed with practice in humans ^1,7–11,14,15^), and substantially reduced the overall learning rate for the new skill ^16^.

In contrast, far less is known about how learning unfolds within individual practice trials, rather than across trials or sessions. The moment-to-moment behavioral dynamics, and their neural correlates, that support online skill acquisition remain poorly characterized. This gap is important because, after only brief practice, real-world performance often shifts from stimulus-driven responding to generative sequence production, in which actions are guided by internal memory representations rather than external cues, unlike in reaction-time paradigms^17^. This rapid transition raises the possibility that the capacity of the underlying skill memory becomes a key constraint on performance.

Evidence from extended training shows that memory capacity increases with practice, in part through chunking, which is the grouping of individual items into larger functional units.^18,19^ With continued training, capacity may increase further as each chunk comes to contain more action elements^20^. These observations suggest that chunking-related transformations may already emerge online during the earliest trials of learning and should be detectable in the fine-grained temporal structure of performance. In early keypress sequence learning, such capacity limits may appear as within-trial periods of high initial skill (HIS) followed by performance declines, consistent with transient information-processing constraints. Within this framework, the number of keypresses typed during an HIS segment (i.e. – HIS segment keypress content) can be treated as an estimate of the effective motor memory capacity available on that trial. If so, HIS segments should be longer for relatively easy, rapidly acquired skills than for slower, more difficult skills^21^, and should lengthen with practice as memory capacity expands^22–24^. By contrast, if within-trial dynamics mainly reflect reactive inhibitory factors such as motor or cognitive fatigue^25,26^, HIS segments should progressively shorten over practice.

At a neural level, theta (θ, 4-8Hz)/gamma (γ, >30Hz) phase amplitude coupling (θ/γ PAC) is a mechanism supportive of memory function^27^. θ/γ PAC refers to the recurrence of identical sequences of γ bursts representing sequential memory items during specific phases of the θ rhythm^28–30^. θ/γ PAC aids in binding individual action elements into coherent sequence representations. Consistently, θ/γ ratios predict behavioral measurements of memory span^31^ and correlate with memory capacity^32^, although this relationship is not specific to any single memory system or task domain^33,34^. β burst oscillatory rhythms have been also linked to memory function^35–37^. Thus, higher θ/γ PAC and β burst oscillatory activity^38–40^ during HIS segments could correlate with motor memory capacity.

Here, we investigated the fine-grained temporal structure of behavior and brain activity that support generative skill acquisition during early ongoing performance using a series of experiments in which human participants learned a sequential keypress skill. We report that (a) segments of high initial skill (HIS) indeed emerged during individual practice trials of early learning across three different skills, (b) the content of HIS segments (number of individual keypresses per HIS segment) was larger when learning fast, easier skills than slow, more difficult ones, (c) both the HIS segment keypress content and chunk content (number of keypresses per chunk) increased with practice in all three tasks, and (d) oscillatory mechanisms implicated in binding actions into memory sequences, theta–gamma phase–amplitude coupling (θ/γ PAC) and beta (β)-band bursts, were elevated during HIS segments, with hippocampal θ/γ PAC predicting HIS segment content.

## Results

In Experiment 1A (N=407), we examined the learning dynamics for generative skills that varied in predicted execution speed due to differences in difficulty (**Figures 1 and S1A-C**). Participants were randomly assigned to practice one of three 6-item keypress sequences over 26 trials (Slow: 4-2-1-3-1-2, Skill Speed Index [SSI] =0.07, N=122; Medium: 3-1-4-2-3-4, SSI=0.41, N=127; Fast: 4-3-2-1-1-1, SSI=0.50, N=158; where “1” is a little finger, “2” is a ring finger, “3” is a middle finger and “4” is an index finger keypress; **Figure 1A**). Each trial comprised 10s of continuous practice followed by 10s of rest^1^, and participants were instructed to perform the sequence as quickly and accurately as possible. Within fewer than six practice trials, most participants could already generate the keypress sequence from memory, indicating that internal representations formed rapidly during early practice across all three skills (Experiment 1B, **Figure S1D-G**).

**Figure 1.**
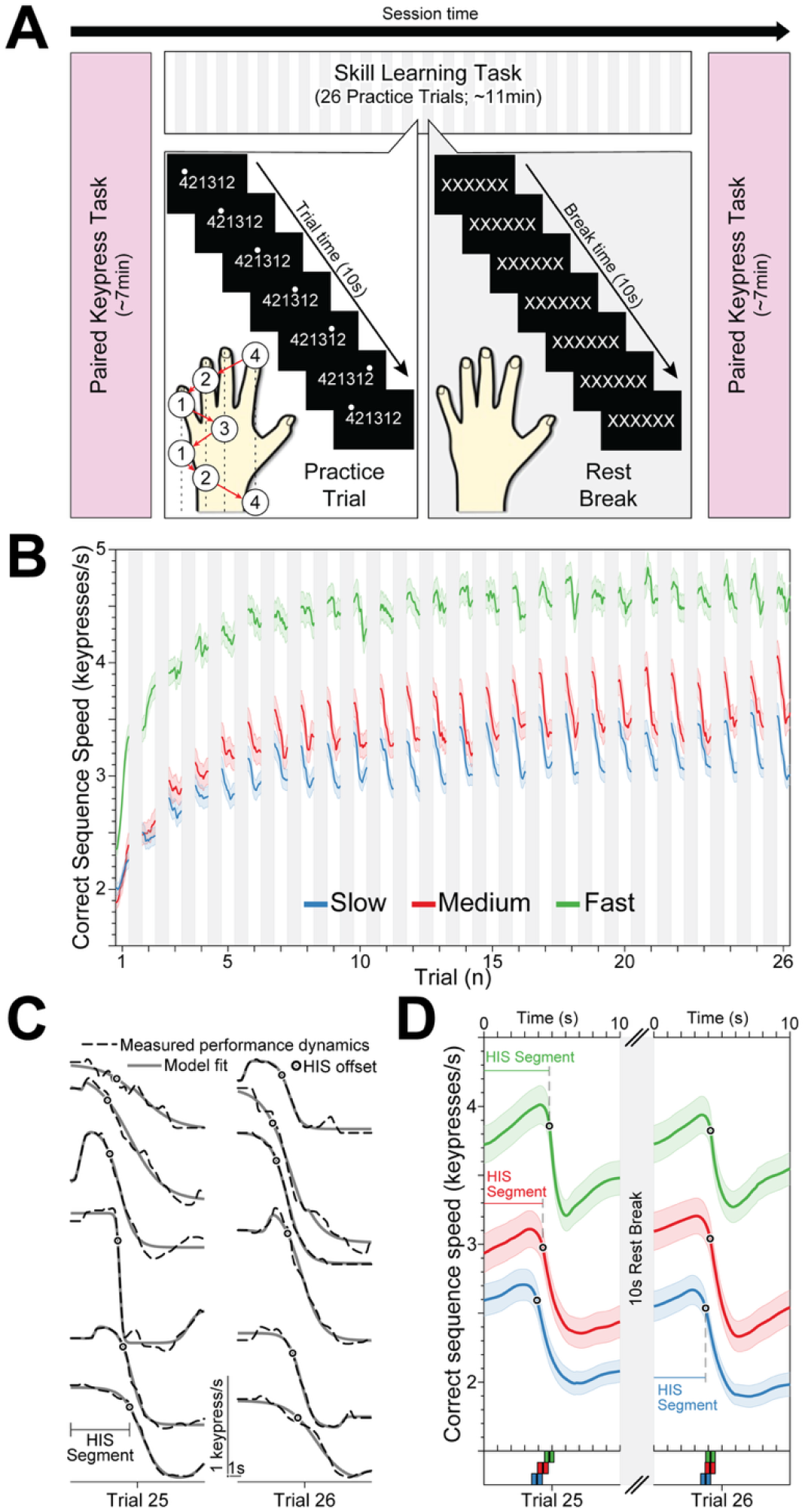
Segments of high initial skill (HIS) emerged with practice across skills. (**A**) We evaluated learning dynamics of three skill sequences of different speed: fast (SSI=0.50), medium (SSI=0.41) and slow (SSI=0.07). Participants (N=407) were randomly assigned to train one of the three skills (slow, medium, or fast) and instructed to perform the task as fast and accurately as possible following an experimental paradigm previously described^1^. Each trial consisted of 10s of practice interleaved with 10s of rest for a total of 26 trials^17^. Prior to and after training, participants also completed the pseudo-randomized paired keypress typing task described in **Figure S1A**. (**B**) Group-level performance curves (correct sequence speed [keypresses/s]^7^, mean±sem) in all participants. Across the three groups, rapid early gains in performance give way to a near-plateau skill level. Arrows indicate the first practice trial in which within-trial skill drops emerge, with these dynamics persisting across subsequent trials to the end of the session. (**C**) We fit a double-sigmoid function to model individual practice trial performance dynamics^41^ (**Figure S2A**). Six representative subject examples of measured performance dynamics (dashed black lines) and model fits (solid gray lines) are shown for trials 25 and 26 (box in **Figure 1B**). Note the high initial skill (HIS) segments, points of maximum deceleration (circles) and following sharp performance declines. Individual lines are arbitrarily shifted vertically for visibility, with scale bars denoting axes scales. (**D**) Group level HIS segment dynamics (trials 25 and 26). Within each skill group, subjects’ models were aligned to the end of the HIS segment (circle) and averaged. Mean and 95% confidence intervals of the end of the HIS segment for each group shown at the bottom.

### Learning leads to stable within-trial performance dynamics characterized by high initial skill (HIS) segments

Practice elicited rapid behavioral gains across all skill groups, with larger improvements in participants training on faster sequences than on slower ones (**Figure 1B**). As trial-by-trial skill gains approached a plateau, performance across all skill groups converged on a shared profile consisting of an initial high, stable or increasing performance segment (high initial skill; HIS segment), followed by a rapid decline and a subsequent partial recovery (**Figure 1C-D**). We parameterized these within-trial fluctuations by fitting the performance time-series from each trial with a double-sigmoid function^41^. The end of the HIS segment was defined as the time-point of maximum deceleration preceding rapid performance drops (**Figure 1C-D**, circles; **Figure S2A**). HIS segments emerged between trials 3 and 5 and persisted through the end of training (**Figure 1D** and **S2B**). Critically, learning was required for HIS segments to emerge since they were absent during practice of pseudo-randomized (non-repeating) keypress sequences executed at speeds comparable to the medium-speed skill (**Figure S3**). These findings suggest that HIS segment dynamics may be shaped by cognitive influences that unfold during learning, such as the gradual accumulation of fatigue during practice, the formation and expansion of skill memory representations, or the planning of upcoming actions.

### HIS segment keypress content scaled with speed of skill practice

Motor fatigue^25,42^ offers one plausible explanation for the within-trial decline in performance observed at the end of HIS segments. Under this account, each keypress incrementally augments fatigue, whereas fatigue dissipation occurs during the intervening keypress transition times (KTTs). Because faster skills involve more keypresses separated by shorter KTTs, they should promote more rapid fatigue accumulation and, consequently, shorter HIS segments. We observed the opposite pattern to that expected under a fatigue account: both HIS segment keypress content and duration increased systematically with skill speed. Mean keypress content was 10.83±0.32 (mean±sem.), 12.63±0.46 and 18.86±0.51 keypresses per HIS segment for the slow, medium and fast skills, respectively (one-way ANOVA, F_2,404_=89.8, *p*<0.001, η^2^=0.31; **Figure 2A**). HIS segment durations were longer for faster skills (fast: 4.72±0.05s) than for slower ones (medium: 4.18±0.07s; slow: 4.14±0.07s; one-way ANOVA, F_2,404_=30.5, *p*<0.001, η^2^=0.13; **Figure S2C**). Together with the absence of HIS segments during practice of pseudo-randomized (non-repeating) sequences (**Figure S3**), the scaling of HIS segment content with skill speed argues against a fatigue-based contribution to HIS segment dynamics.

**Figure 2.**
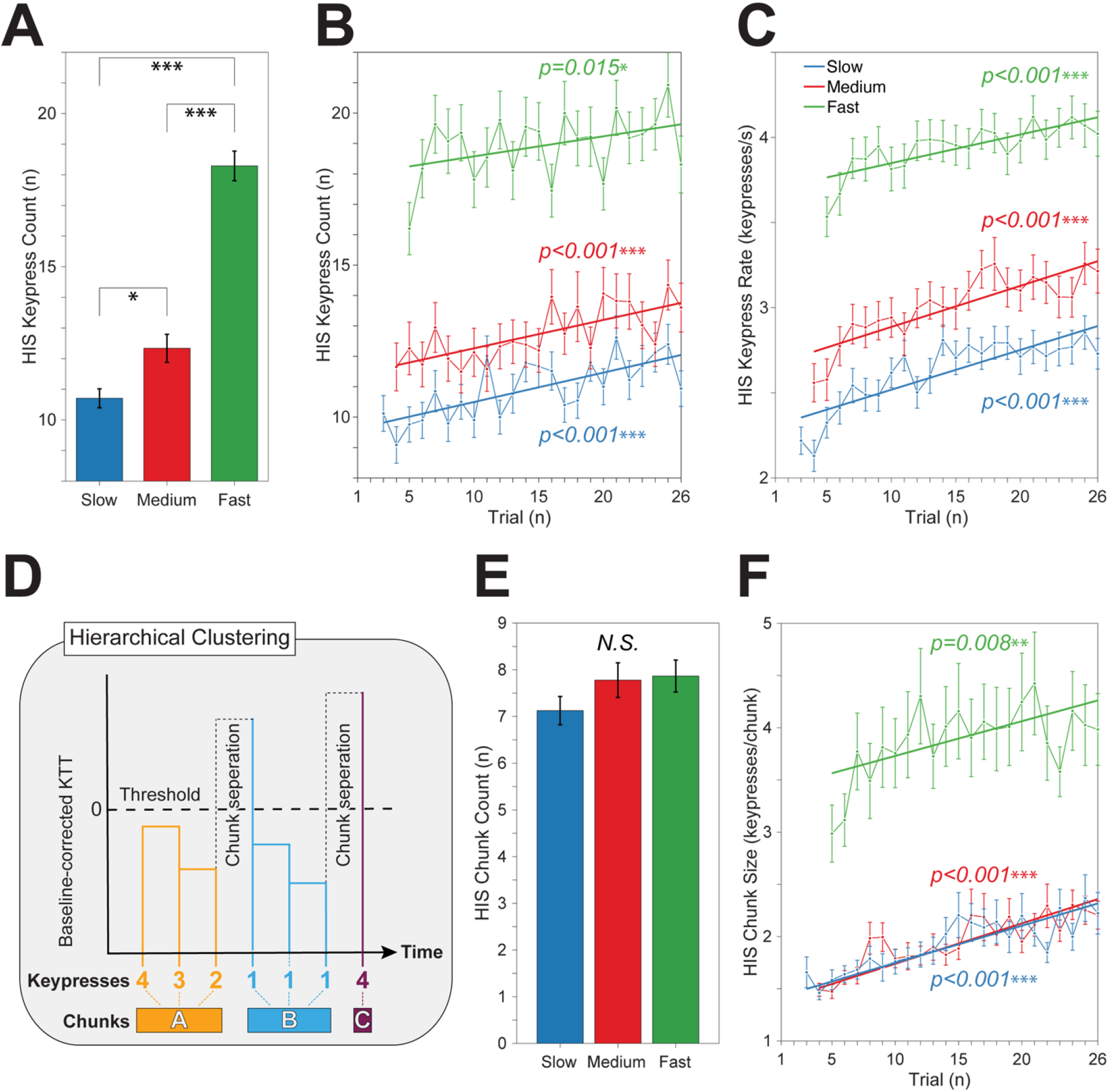
HIS segment keypress content increased with practice. (**A**) HIS segment keypress content ranged between 10 and 18 keypresses and increased with skill speed. Asterisks indicate statistically significant differences during post-hoc comparisons (*p < 0.05, **p < 0.01, ***p < 0.001). (**B**) HIS segment keypress content increased progressively with practice in all skill groups (keypresses, mean±sem; BHFDR-corrected LMMs). Solid lines show the linear-fitted model for each group. Asterisks indicate statistically significant progressive increase in HIS keypress content with training. (**C**) The practice-dependent increase in HIS segment keypress content was not simply driven by changing HIS segment durations, as keypress rates— controlling for this factor—also increased progressively with practice across all skill groups (Slow: F_1,2629_=143.0, *p*<0.001, 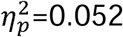; Medium: F_1,2660_=121.2, *p*<0.001, =0.044; Fast: F_1,3199_=43.8, *t*_3199_=6.62, *p*<0.001, 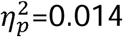). (**D**) Hierarchical clustering was applied to within-individual, KTT_pairs_-corrected KTT_seq_ values (KTT_seq_ − KTT_pairs_; see **Methods**). Chunk boundaries were defined at the point where KTT_seq_ exceeded KTT_pairs_ (threshold = 0), such that longer-than-KTT_pairs_ transitions marked separations between clusters (chunks). (**E**) The mean number of chunks within HIS segments ranged between 7 and 8 for all skills. (**F**) During HIS segments, chunk content (keypresses per chunk) increased progressively with practice in all skill groups (keypresses per chunk; BHFDR-corrected LMMs).

### Training-Related Expansion of HIS Segment Content Persists After the Skill Plateau Was Reached

Alternatively, HIS segment dynamics may reflect the formation and expansion of the skill memory over the course of practice. Across all skill groups, the number of keypresses within HIS segments (i.e. - HIS keypress content) increased with training (BHFDR-corrected linear mixed-effects models [LMMs]: Slow, F_1,2633_=32.0, p<0.001, 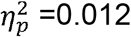 Medium, F_1,2663_=21.9, p<0.001, 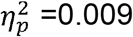 Fast, F_1,3201_=5.90, p=0.015,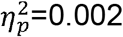 **Figures 2B-C**).

To determine whether the expansion of HIS segment content persisted after performance stabilized, we repeated the analysis using only the plateau phase of the learning curves (trials 6–26). During this phase, the number of keypresses executed within HIS segments continued to increase in the Slow and Medium groups (Slow, F_1,2294_=16.03, p<0.001, 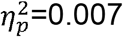 Medium, F_1,2431_=18.9, p<0.001, 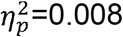), but not in the Fast group (F_1,3046_=1.65, p=0.198,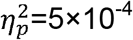). By contrast, HIS-segment keypress rates (Slow: F_1,2291_=45.2, p<0.001; Medium: F_1,2430_=58.4, p<0.001; Fast: F_1,3044_=24.4, p<0.001) and skill speed (**Figure S4**) continued to increase in all groups. These results indicate that HIS segments continue to expand even after overall behavioral performance has plateaued.

### Training-dependent enrichment of HIS segments is contingent upon the expansion of motor chunk content

Next, we asked whether changes in chunking organization could account for the training-dependent expansion of HIS segment content^24^. We used hierarchical clustering to identify chunk boundaries within HIS segments (**Figure 2D**). For each participant, a boundary was defined wherever the duration of a keypress transition within the sequence (KTT_seq_) exceeded the corresponding transition duration measured outside the skill learning context (KTT_pairs_)^43–46^. This procedure revealed a clear chunking structure in HIS segments across all skill groups.

The overall number of chunks per HIS segment was comparable across skills (slow: 7.13±0.30; medium: 7.78±0.37; fast: 7.87±0.34 chunks/HIS; one-way ANOVA, F_2,404_=1.33, *p*=0.27, η^2^=0.003; **Figure 2E**). In contrast, the number of keypresses per chunk was greater for the fast skill than for the medium or slow skills (Slow: 1.93±0.10, Medium: 1.95±0.08, and Fast: 3.84±0.26 keypresses per chunk, one-way ANOVA, F_2,404_=36.5, *p*<0.001, η^2^=0.12). Critically, the number of keypresses per chunk increased progressively with practice in all skill groups (Slow: F_1,2630_=72.1, *p*<0.001, 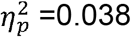, Medium: F_1,2662_=81.5, *p*<0.001, 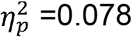, Fast: F_1,3200_=9.03, *p*=0.003, 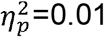, **Figure 2F**) and this effect persisted when analyses were restricted to the performance plateau (trials 6–26; Slow: F_1,2292_=39.2, *p*<0.001; Medium: F_1,2432_=45.7, *p*<0.001; Fast: F_1,3045_=3.98, *p*=0.046). Thus, training also expanded the keypress content of individual chunks across skills. Taken together, these behavioral findings raise the possibility that HIS dynamics reflect a pre-planning effect or alternatively, expansion of a skill-memory buffer, rather than the accumulation of motor fatigue.

### HIS Segment Dynamics Are Not Explained by a Pre-Planning Advantage at Trial Onset

We next tested whether HIS dynamics could be explained by known pre-planning effects. Prior work on stimulus-driven sequential behavior indicates that pre-planning increases the execution speed of the initial 3 to 4 actions relative to subsequent responses within a given trial^47–53^. Thus, we quantified a putative pre-planning effect on initial performance within each trial by comparing the execution speed of the first three keypresses in the first sequence (which fall within the planning horizon and thus should be performed faster in the case of a pre-planning benefit) with the same three keypresses in the subsequent sequence (which fall outside the horizon). Consistent with our model-based analysis of within-trial performance dynamics (see initial upward trend in HIS segments in **Figure 1C-D**), participants either maintained a constant speed or accelerated across the first two sequence iterations, both before HIS segments emerged (**Figure S2D, E)** and after their emergence (**Figure S2D, F**). Thus, we found no evidence for a pre-planning advantage in any of the three skill groups.

### Increased θ/γ PAC and β band bursting rates during HIS segments of early learning

In **Experiment 2** (N=32), participants learned a medium-speed sequence in a magnetoencephalography (MEG) setting^1^. The goals were (i) to test the reproducibility of the behavioral effects described above and (ii) to identify physiological markers of memory expansion during HIS segments. Behaviorally, the results closely matched those from Experiment 1. Performance time courses showed the emergence of HIS segments beginning at trial 5 and persisting through the end of training (**Figure S5A-B**). The HIS segments keypress content (14.07±0.91) increased with practice (LMM: F_1,985_=9.00, *p*=0.003) while HIS segment duration remained stable (F_1,1020_=1.21, *p*=0.27, 4.10±0.11, ≈4.0 s). Thus, training-dependent expansion of keypress content within HIS segments was robustly replicated across two independent experiments.

Next, we examined θ/γ phase–amplitude coupling (PAC) ^27,54,55^. Whole-brain source-space θ/γ PAC was significantly higher during HIS segments (trials 5–36) than during the corresponding time window in pre-HIS trials (trials 1-4, paired *t*-test, *t*_31_=-2.95, *p*=0.006, **Figure 3A**). Parcellated source-space analyses revealed θ/γ PAC during HIS segments to be significantly elevated relative to time-shifted surrogates in contralateral hippocampus, insula, and orbitofrontal cortex, bilateral ventromedial prefrontal cortex, and ipsilateral parieto-occipital regions (cluster-forming threshold *t*>2.74 and distance<20mm, number of permutations=2000, *p*<0.01, **Figure 3B, Table S1**). In these regions, θ/γ PAC was greater during HIS segments (0–4 s of each practice period) than during post-HIS intervals of the same trials, whether defined as 4–10 s (**Figure 3C**) or as a duration-matched 4–8 s window (**Figure S6A**). Critically, hippocampal θ/γ PAC strongly predicted HIS segment content (keypresses/HIS segment; LMM, F_1,986_ =13.5, *p*<0.001; **Figure 3D**) and tracked skill level across practice (LMM, F_1,1061_=6.21, *p*=0.013; **Figure S6B**). Consistent with these findings, an exploratory trial-by-trial analysis showed that θ/γ phase-amplitude coupling (PAC) within HIS segments increased progressively with practice (**Figure S6C-D**).

**Figure 3.**
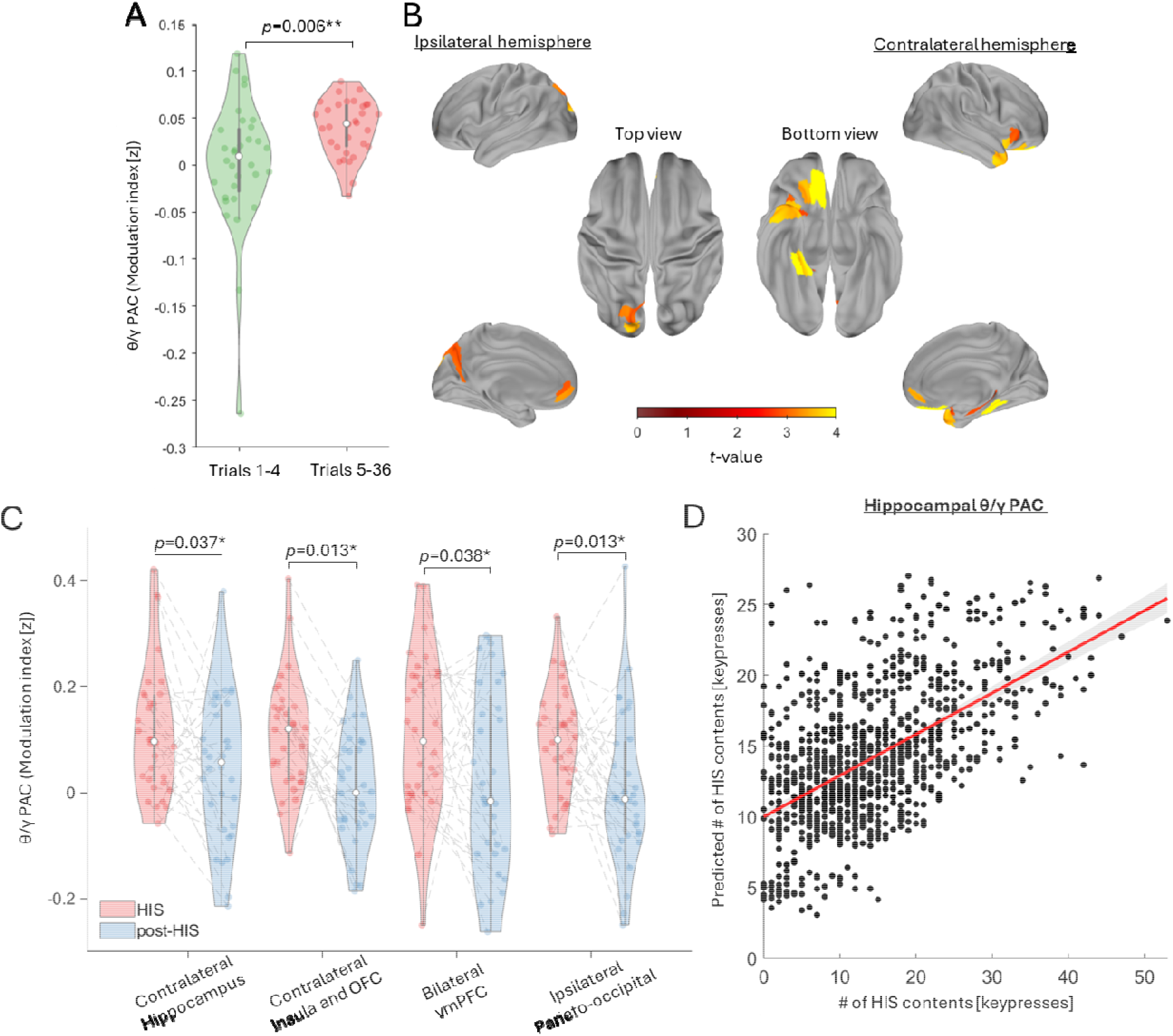
θ/γ PAC, HIS segment content and overall skill. (**A**) Whole-brain source-space parcellated MEG θ/γ PAC was significantly higher during HIS segments (trials 5-36) than in the same time window of trials preceding the appearance of HIS segments (trials 1-4). Removal of two outliers did not modify the outcome (paired t-test, *t*_29_=-2.56, *p*=0.016). (**B**) During HIS segments (trials 5-36), θ/γ PAC strength was significantly higher in contralateral hippocampus, insula and orbitofrontal cortex (OFC), bilateral ventromedial prefrontal cortex (vmPFC), and ipsilateral parieto-occipital regions (relative to time-shifted 200 surrogates^114^; **Table S1**). (**C**) θ/γ PAC during HIS segments (0-4 s of each practice period) was significantly higher than following HIS segments (4-10s of each practice period) in all regions identified in **Figure 3B** (paired *t*-test, BFDR corrected; **Figure S6A**). (**D**) Out of these regions, only hippocampal θ/γ PAC predicted HIS segment content (R^2^=0.28) as well as overall skill (**Figure S6B**).

Examination of oscillatory bursting across frequency bands revealed selective effects in the β range. β-band burst rates were higher during HIS segments than during post-HIS segments within the same trials (whole-brain MEG sensor space; **Figure 4A**), with no corresponding differences in other bands. β burst rates were also greater in trials containing HIS segments (trials 5–36) than in the equivalent time window of trials 1–4, before HIS segments appeared (paired *t*-test, *t*_31_=−3.71, *p*<0.001; **Figure 4B**). Source-space parcellation showed significantly elevated β burst rates during HIS segments relative to post-HIS intervals in bilateral insula and frontal, temporal, and anterior cingulate cortices (cluster-forming threshold *t*>3.37, distance<20 mm, 2000 permutations, *p*<0.01; **Figure 4C, Table S2**).

**Figure 4.**
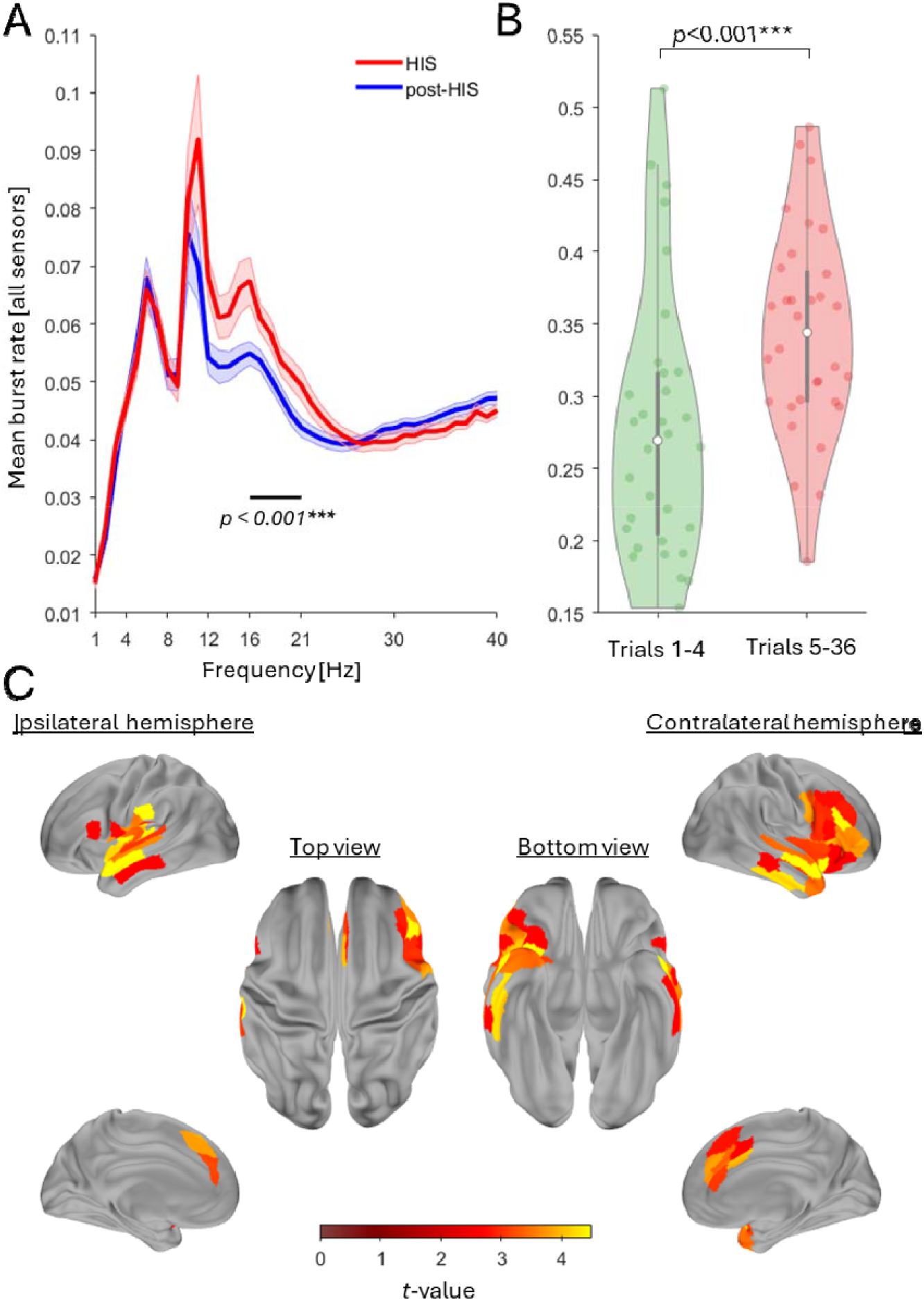
Beta burst activity during HIS segments. (**A**) Broad-band whole-brain sensor-space MEG oscillatory burst rates^38^ during HIS- and post-HIS segments (trial 5-36; HIS segment was determined in each trial and each participant). Note the significantly higher burst activity in the 16-21 Hz range during HIS relative to the post-HIS segments. β band bursts during HIS segments displayed higher power, higher frequency, and longer duration compared to bursts observed following HIS segments (**Figure S7**) (**B**) β burst rates in the 0-4 s window corresponding to the HIS segment were significantly higher in trials containing HIS segments (trials 5–36) than in trials preceding the emergence of HIS dynamics. (**C**) Bilateral insula and frontal, temporal and anterior cingulate cortices were the main contributing regions to the higher source-space parcellated MEG β burst activity during HIS segments (**Table S2**).

## DISCUSSION

Here, we show that generative skill learning is marked by the emergence of a reproducible within-trial performance pattern consisting of high initial skill (HIS) segments followed immediately by rapid slowing. HIS segments contained more sequential keypresses for faster than for slower skills, and their keypress content increased progressively with training, indicating that increasingly larger action sequences became integrated into single performance chunks. Importantly, HIS segments emerged only with learning, as they were absent during practice of pseudo-randomized, non-repeating keypress sequences of a comparable speed. These properties are difficult to reconcile with an account based primarily on motor fatigue or known pre-planning advantage effects. Instead, they point to the formation and expansion of skill-memory representations over practice. Consistent with this interpretation, neural activity during HIS segments was characterized by elevated θ/γ phase–amplitude coupling (PAC) and β-band bursts across memory-related neocortical and medial temporal regions, with hippocampal θ/γ PAC strongly predicting HIS segment content.

### HIS segment dynamics are not explained by motor fatigue

Practice led to prominent initial performance improvements followed by a phase in which average skill stabilized. We found that segments of high initial skill interrupted by skill drops emerged after the initial training trials (practice trials 3 to 5 depending on skill speed) and persisted along the plateau period in all subjects and groups. We and others have attributed these performance drops to fatigue accumulating with continued practice^1,56^, particularly during the execution of fast motor tasks^56,57^. Under this scenario, each keypress results in some amount of fatigue that is recovered from during the “resting” time between keypresses (KTT)^57,58^. We reasoned that if this were the case, HIS segment content should be smaller when learning fast skills (containing more fatigue-inducing keypresses combined with shorter recovery time intervals) than for slower skills (fewer keypresses combined with longer recovery time intervals). Contrary to this prediction, we found that the keypress content of HIS segments was larger for faster skills compared to slower ones, ranging between 10 and 18 keypresses (**Figure 2A**).

One would also expect that the accumulation of motor fatigue with practice^25,26,57^ would result in progressively slower typing speed and/or earlier skill drops, leading to shorter HIS segment durations with diminished keypress content. Contrary to these predictions, we observed a training-dependent expansion of HIS segment keypress content and keypress rates across all skills tested (**Figure 2B-C**). These findings are inconsistent with a fatigue-based explanation of HIS segment content, aligning instead with reports that motor fatigue requires practice durations longer (10–60 min) than those used in this study (<9 min)^59,60^. Thus, motor fatigue does not appear to play a pivotal role in shaping HIS segment dynamics during the early stages of mastering a generative sequential skill.

### HIS segment capacity extends beyond established planning horizon limits

One possible interpretation is that HIS segment dynamics reflect advanced planning. However, this view is difficult to reconcile with evidence from diverse motor tasks and modalities showing that the planning horizon is typically restricted to fewer actions^47–53,61–65^. In skilled sequential performance, advanced planning extends roughly 1–2 seconds into the future^47^; coarticulation effects likewise tend to influence only the next 1–3 actions^50^; and perturbation effects generally dissipate within 1–2 subsequent actions^66^. Taken together, these findings indicate that, particularly in non-expert performance such as that studied here, advance planning usually spans approximately 2–4 individual actions^67,68^. By contrast, the HIS segments observed in the present study comprised 10–18 keypresses (**Figure 2A**), far exceeding the planning horizon reported in the literature^61^. Consistently, we found no selective speed advantage for the first 3 keypresses of the initial sequence within a trial relative to the corresponding keypresses in the subsequent sequence, as would be expected if advanced planning contributed substantially to HIS dynamics. Notably, this absence of a planning-related advantage was evident throughout training and both before and after the emergence of HIS segments (**Figure S2D–F**).

Nevertheless, it remains theoretically possible that, under some circumstances, the planning horizon could extend over longer strings of actions than those typically reported in the literature^47–53,61–65^. Such expanded planning has been documented in highly practiced domains: expert typists can prepare approximately 1–2 words (8–15 letters) ^69^, concert pianists up to 1–2 measures (12–20 notes) ^70^, and sequence learners as many as 8–12 items once the sequence has been extensively practiced and consolidated^71^. Importantly, however, these findings come from contexts involving substantial training and, often, years of skill acquisition. While a contribution of longer-range planning cannot be fully excluded, the literature suggests that more extensive practice than the 5–7 brief 10-second trials administered here would be required to support planning across the full 10–18 keypresses of the HIS segments.

### HIS segment dynamics inform on motor memory capacity during early skill learning

While HIS segment dynamics are not well explained by fatigue or pre-planning accounts, they may index the emergence and deployment of learned internal representations capable of supporting prospective control^47–53^. The observation that HIS segments comprised 10–18 keypresses (**Figure 2A**), substantially more than typical capacity limits reported in other memory domains^72–74^, argues against the view that individual keypresses remain the basic units of sequential skill memory with continued practice. Instead, a more plausible account is that practice transforms these elementary actions into higher-order memory units, or chunks, of keypresses. This possibility is consistent with classic accounts of motor skill learning, in which repeated actions are progressively grouped into chunks that reduce the demands of sequential control and memory storage when sequence length exceeds capacity limits^24^. We therefore asked whether the structure of HIS segments could be explained by changes in chunking organization. To identify chunk boundaries within each skill sequence, we applied hierarchical clustering to keypress transition times (KTTs), defining chunks as action groups bounded by relatively long transitions ^44–46 44,75,76^. Strikingly, this analysis revealed that HIS segments contained a highly similar number of chunks across skill speeds, approximately seven (**Figure 2E**), a value that matches canonical estimates of memory capacity in other domains, including working memory (3–7 items) ^72–74,77^. Together, these findings suggest that the functional units of sequential skill memory during early learning may be action chunks rather than individual keypresses.

Next, we examined how practice influenced chunking organization. Across all skill levels, the number of keypresses per chunk expanded with training rather than contracting (**Figure 2F**), a pattern also inconsistent with a fatigue-based account. Notably, this content continued to expand even during the plateau phase of the learning curves, despite overall performance speed reaching near-ceiling levels (**Figure S2G**). An interesting finding was the relatively stable HIS segment duration of approximately 4 seconds across training. It is possible this transition point represents an upper temporal limit over which nascent optimal performance states can be sustained^78^, possibly related to limits in attentional stability^79,80^, temporal properties of skill memory retrieval^81^, or stability of emergent hippocampo-neocortical rhythms supporting skill performance^82^.

Altogether, these results are consistent with HIS segment dynamics reflecting expansion of the skill memory capacity in the course of early learning^83^. This interpretation accords with contemporary views of memory, and hippocampal memory in particular, not as a passive storage system, but as a predictive architecture that supports the simulation of future states and the guidance of goal-directed behavior^81,84^. In this view, memory is not simply a record of past experience; it provides a generative model that can be used to anticipate, evaluate, and organize possible future actions. Processes such as neural replay^85^ may contribute to this function by linking past experience to prospective control, thereby supporting planning, decision-making, recollection, and consolidation. In this framework, the progressive expansion of HIS segments may reflect the increasing capacity of learned memory representations to prospectively structure longer stretches of behavior during skill acquisition^81,84^.

### Larger θ/γ PAC and β-band bursting during HIS segments

How might the expansion of memory capacity during early learning be supported at a neural level? Theta–gamma phase–amplitude coupling (θ/γ PAC) has been implicated in binding individual actions into coherent sequences^32,86–94^ and in the active maintenance of information within other memory types^27,54,55^. Our observation of elevated θ/γ PAC during HIS segments suggests that this coupling may facilitate the integration of individual finger representations, reflected in low frequency γ bursts^95^, into precise temporal and spatial positions within slower θ oscillations (i.e. precession), thereby encoding the identity of emerging sequences and chunks^32,86–94^. This binding mechanism, operating during motor execution, is likely reinforced during rapid consolidation periods that develop across rest intervals interspersed with practice^1^. Contributors to rapid consolidation during these rest periods of early skill learning include wakeful neural replay^7,16^ and contextualization of individual sequence action representations^17^.

Elevated θ/γ PAC during HIS segments was observed across a distributed network encompassing memory-related and sensorimotor integration regions (including the hippocampus, insula, orbitofrontal, prefrontal cortex, and regions within parieto-occipital and intraparietal sulci). In this context, the hippocampus emerged as a key node, as θ/γ PAC in this region strongly predicted HIS segment content (**Figure 3D**) and actual skill (**Figure S6B**). This observation aligns with evidence implicating hippocampo– neocortical neural replay^1,7–11^ and hippocampal ripples, established biomarkers of replay events,^96^ in supporting micro-offline performance gains during early phases of generative skill acquisition^14–16^. Altogether, these findings converge with the well-documented role of the hippocampus in supporting skill memory formation^5,7,8^.

Finally, we evaluated broadband neural oscillatory features associated with HIS segment content. The only frequency band differentiating transient high-power bursts during HIS versus post-HIS segments was β (16-21 Hz), which revealed elevated burst rates during HIS segments particularly in the insula and frontal, temporal and anterior cingulate cortices. Furthermore, whole-brain β burst rates were higher in trials after HIS segments had become clearly defined (trials 5-36) than in trials preceding the emergence of HIS segments (trials 1-4). These results are in line with previous work showing that high β oscillatory activity ^97^, organized predominantly in bursts^39,98,99^, supports skill memory function. Thus, higher θ/γ PAC and β burst oscillatory activity during HIS segments constitute potential mechanisms supporting the expansion of memory capacity and content during early learning of generative sequential skills.

### Limitations and future directions

In the present study we evaluated chunking of generative, self-paced skill sequences using keypress timing patterns, which were generalizable across four different skill sequences and two experimental platforms (online and in-lab). It would be important for future in-lab investigations to evaluate chunking patterns of finger kinematics that evolve with learning, to determine if these patterns are consistent with the present keypress timing observations. In the present study, we assessed neural substrates of HIS segment content for only one of the skill sequences, which was performed at a medium speed level. Future work could expand this investigation to determine if these findings generalize across different skill sequences performed at lower or higher speeds, as well. Finally, we cannot exclude the possibility that performance-inhibiting factors like motor fatigue may exert a greater influence on HIS segment dynamics over much longer or more effortful training protocols than those examined here.

In summary, our findings suggest that generative skill learning may be contingent upon a gradual augmentation of skill memory capacity, as evidenced by the content of HIS segments. Notably, this increase persists even after the overall skill speed performance reached a plateau. Elevated θ/γ PAC and β-band burst rates across memory-related neocortical and medial temporal regions, particularly hippocampal θ/γ PAC, emerged as candidate neural mechanisms supporting this expansion and skill gains.

## STAR★Methods

### Resource Availability

Further information and requests for resources should be directed to and will be fulfilled by the Lead Contact, Leonardo G. Cohen

### Materials availability

This study did not generate new materials.

### Data and Code availabilit

- All de-identified and permanently unlinked from all personal identifiable information (PII) data are publicly available on the OpenNeuro platform. (https://openneuro.org/datasets/ds006502/versions/1.0.0).
- All custom analysis code will be available in a publicly accessible repository hosted on GitHub.

## EXPERIMENTAL MODEL AND STUDY PARTICIPANT DETAILS

### Participants

Right-handed participants performed a well-established sequential motor learning task^100,101^ consisting of repetitive typing of keypress sequences with their non-dominant, left hand. We initially collected a very large set of paired keypress transition data (N=2152, 1043 women, mean age 28.1±0.12) utilizing the *Connect* crowd-sourcing platform (CloudResearch, NY, USA). This independent dataset was used to generate the Skill Speed Index (SSI) crucial for sequence selection to assess skills at different predicted speeds. The primary study consisted of two experiments: Experiment 1 (N=407, 199 women, mean age 28.1±0.26) was also conducted online utilizing the *Connect* platform, while Experiment 2 (N=32, 16 women, mean age 26.6±0.87) was carried out at the National Institutes of Health (NIH) Clinical Center. The study was approved by the NIH Institutional Review Board (IRB). All online participants (i.e. – the preliminary paired keypress transition dataset and Experiment 1 from the primary study) completed a digital acknowledgement of participation, and in-person participants (Experiment 2) completed the informed consent process before engaging in study procedures.

## METHOD DETAILS

### Task

We first collected a large dataset of independent keypress transition times (KTTs) from healthy volunteers (N=2152) performing all 16 possible two-finger transitions using the little, ring, middle, and index fingers of their non-dominant (left) hand on a standard keyboard (**Figure S1A**). On each trial, participants saw a pair of numeric cues (e.g., “24”) indicating the required finger pair (1 = little, 2 = ring, 3 = middle, 4 = index) and were instructed to respond as quickly and accurately as possible. Individual numbers were replaced with “X” as the corresponding keypresses were executed. Incorrect responses triggered a repetition of the trial until a correct response was registered. The inter-trial interval was uniformly jittered between 200–500 ms, and each of the 16 possible transitions were sampled five times in pseudorandom order.

In Experiment 1A, we evaluated learning dynamics for three skill sequences with different predicted speed profiles (Slow, n=122 [28.4±0.5 years-old]; Medium, n=127 [28.8±0.47 years-old]; Fast, n=158 [28.1±0.1 years-old]; **Figure S1C**). Participants completed a single online session using their own personal computer. The session included the pseudorandomized paired keypress transition task completed before and after a motor skill learning task where they repeatedly practiced the same short 6-item keypress sequence (**Figure 1A**)^102,103^. The purpose of the paired keypress transition task was two-fold: (1) to familiarize all participants with the stimulus-response mapping (e.g. – “4” corresponds to an index finger keypress) and (2) to provide baseline transition data for the sequence chunking analysis pipeline.

Skill training comprised twenty-six 10-second trials, which alternated with 10-second of rest breaks, adhering closely to methodological recommendations for minimizing reactive inhibition^56^. Participants were instructed to perform the displayed sequence as fast and accurately as possible, repeating it continuously during each practice interval, and were randomly assigned to one of the three skill speed groups (Slow: 4-2-1-3-1-2; Medium: 3-1-4-2-3-4; Fast: 4-3-2-1-1-1; **Figure S1C**; where 1, 2, 3, and 4 corresponded to the little, ring, middle and index fingers, respectively; **Figure 1A**).

The sequence remained visible throughout practice. A small dot appeared after each keypress to indicate the participant’s current position within the sequence, and after each full sequence iteration the dot returned to the first item. No explicit performance feedback was provided. During rest intervals, the practice sequence was replaced with “XXXXXX” and no dot was displayed. Throughout the experiment, keypress timing and finger identity were recorded, and skill was quantified as the speed of correctly executed sequences (keypresses per second)^1^.

In Experiment 1B, we assessed the memorability of the three sequence skills in an independent dataset (Fast: N=63 [28.1±0.1 years-old]; Medium: N=69 [26.7±0.1 years-old]; Slow: N=71 [27.2±0.1 years-old]). Participants completed 26 practice trials with alternating task demands. On odd-numbered trials (1, 3, …, 25), the sequence remained visible, allowing visually guided practice of sequential actions. On even-numbered trials (2, 4, …, 26), participants executed the sequence from memory, without the target sequence on a display. Numeric sequence stimuli were replaced with “?”s and the same position feedback (i.e. - the jumping dot) was provided (**Figure S1D**). Memorability was operationalized as the proportion of correctly generated sequences (e.g. sequence accuracy) during each memory-guided trial.

Experiment 2 was carried out in-person in the NIH magnetoencephalography (MEG) lab^104^. Here, subjects practiced a medium-speed sequential skill (4-1-3-2-4 where 1, 2, 3, and 4 corresponded to the little, ring, middle and index fingers, respectively) over thirty-six 10-second trials, which alternated with 10-second rest breaks. Subjects used an MEG-compatible response pad (Cedrus LS-LINE, Cedrus Corp.) to record keypresses.

### Behavioral Data Analysis

#### Online Data Collection Quality Control

Participants were recruited through the *Connect* (CloudResearch)^105^ crowd-sourcing platform, which incorporates both respondent vetting and technical fraud-prevention procedures intended to improve data quality. *Connect* uses Sentry® (CloudResearch’s proprietary screening system) during participant registration to assess several indicators relevant to compliant participation including: attention, honesty, language comprehension, and the ability to follow instructions. Sentry® also flags response patterns associated with low-quality or fraudulent participation, including use of survey translation tools, failure to follow instructions, copy-pasted responses, automated responding, and poor-quality open-ended answers. In addition, *Connect* applies technical checks to reduce duplicate or fraudulent participation, including verification of single-account use, consistency between the participant’s IP address and reported U.S. location, uniqueness of payment accounts, and detection of devices shared across multiple participant accounts.

Furthermore, compliance was specifically evaluated in our study using both post-experiment self-report questionnaires and behavioral indices extracted from the recorded keypress data. Participants were excluded if they reported using their right hand, indicated that they thought they had to execute the sequence only once per practice trial, or showed absence of keypresses lasting 5 s or longer in more than four practice trials. On this basis, 290 (out of 899) total participants were excluded from analyses.

#### Skill Speed Index (SSI)

The Skill Speed Index (SSI) was derived from z-standardized keypress transition times (KTT_pairs_) previously collected from a large independent dataset (N=2152) using a paired keypress task (**Figure S1A-B**). This task repeatedly sampled all possible paired keypress transitions (16 total possibilities for the four fingers). Positive z-score values corresponded to relatively fast transitions (e.g., 4–4, 3–3, 2–3), whereas negative values reflected slower transitions (e.g., 1–3, 4–2, 2–4). SSIs were calculated for all possible 6-item sequences by summing the corresponding KTT z-scores and subsequently applying min–max normalization across the entire set (**Figure S1C**).

Three sequences, containing at least one keypress for each finger, with different SSIs but similar transition entropy values were selected from this set (*Slow*: SSI = 0.07, H = 2.58; *Medium*: SSI = 0.41, H = 2.58; *Fast*: SSI = 0.50, H = 2.25; **Figure S1C**) and utilized in Experiment 1 of this study.

#### Skill Measurement

We quantified performance by measuring the time required to complete each correct sequence iteration and by calculating the rate of correct keypresses per second (keypresses/s). A keypress was considered correct if it matched any circular shift of the target sequence. Skill values derived from individual correct keypresses were then interpolated and extrapolated to yield a continuous, millisecond-resolution measure of skill across the entire practice interval^7^. We excluded the initial reaction time (from practice trial onset to the first keypress) and any transitions containing errors^106^.

#### Within-trial Skill Dynamics

First, we used a regression model to identify the onset of practice trials yielding within-trial performance declines characterized by high initial skill (HIS) segments (**Figure 1B-D**):

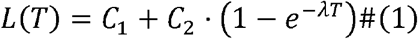

where *L*(*T*) represented the group averaged linear within-trial skill change on practice trial, *T, C*_1_ and *C*_2_ controlled the initial performance and asymptote of the exponential term, and λ controlled the rate of change. Parameters *C*_1_ ∈ [0,5], *C*_2_ ∈ [-5,5], and λ ∈ [0,2] were estimated using a constrained nonlinear least-squares method (*lsqcurvefit*.*m*, trust-region-reflective algorithm). The point at which *L*(*T*) decreased below zero was used to define HIS emergence for each skill.

Next, we identified the end of each HIS segment for trials revealing the within-trial skill drop by fitting a double-sigmoidal function *S*(*t*) to the min–max normalized skill time series. This model comprised three components: a plateau skill level, P, within-trial gain dynamics, *G*(*t*), and within-trial drop dynamics, *D*(*t*):

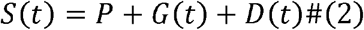

where,

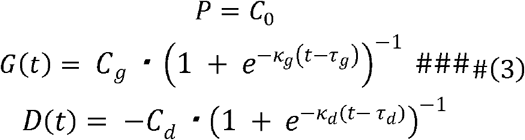

Here, *C*_0,*g,d*_ controlled the magnitude of each component, *κ*_*g,d*_ the activation rate, and *τ*_*g,d*_ the activation latency of the positive and negative sigmoidal functions. All parameters were constrained within the following bounds: *C*_0,*g,d*_ ∈ [0,1], _g,d_ ∈ [0,0.5], and *κ*_*g,d*_ E[0,10000].

Parameters were optimized using 600 random initializations per trial: 300 for the single-sigmoid model and 300 for the double-sigmoid model (MATLAB lsqcurvefit, trust-region-reflective algorithm). For each trial, the parameter set with the lowest Bayesian information criterion (BIC) was retained, providing a principled trade-off between goodness of fit and model complexity. Consistent with the predominantly non-monotonic within-trial performance dynamics, the single-sigmoid model was selected as the best-fitting model in only 1.13% of trials, whereas the double-sigmoid model was favored in 98.87%.

This modeling framework effectively isolated within-trial skill decay *D*(*t*) from learning-related gains. The end of each HIS segment was defined as the time of maximal deceleration in *D*(*t*) (i.e., the second derivative peak; **Figure 1C-D** and **Figure S2A**). We then quantified HIS segment content (keypress #, keypress rates) changes within and across-individuals.

#### Chunking organization

To identify chunk boundaries within learned motor sequences, we applied hierarchical clustering to individual keypress transition times (KTTs) during HIS segments^43^. Boundaries were defined as transitions that were slower during sequential execution (KTT_seq_) than in the corresponding non-sequential context (KTT_pairs_), indicating natural segmentation points in performance. By contrast, transitions within a chunk were characterized by KTT_seq_ values that were equal to or shorter than KTT_pairs_, consistent with prior evidence that transition times are reduced within well-learned chunks ^44–46^. A dendrogram was created using the within-individual, absolute KTT_pairs_-corrected KTT_seq_ values to group temporally adjacent transitions (**Figure 2D**). The dendrogram was then cut at a threshold of zero, such that any KTT_seq_ longer than its corresponding KTT_pair_ defined a chunk boundary. We then quantified the number of chunks, average chunk keypress content, and chunk rate within each HIS segment.

#### MRI acquisition

Structural MRI scanning was performed on a 3T MRI scanner (GE Excite HDxt and Siemens Skyra) with a 32-channel head coil. High-resolution, T1-weighted anatomical images (1 mm^3^ isotropic MPRAGE sequence) were acquired for each participant in the in-lab experiment to perform surface-based cortical dipole estimation from MEG signals.

#### MEG acquisition

Continuous MEG was recorded at a sampling frequency of 600 Hz with a CTF 275 MEG system (CTF Systems, Inc., Canada) while participants were seated in an upright position. The system is composed of a whole head array of 275 radial 1st-order gradiometer / SQUID channels housed in a magnetically shielded room (Vacuumschmelze, Germany). Three of the gradiometers were malfunctioning and were not used for all recordings, resulting in 272 total MEG channels. Background noise was removed using synthetic 3rd order gradient balancing. Temporal alignment of the behavioral and MEG data was carried out using a TTL trigger signal. Head position indicator points were monitored before and after the recording in the scanner coordinate space using head localization coils placed on the nasion, left and right pre-auricular locations of the participant’s head. These coil positions were also recorded in the subject’s T1-MRI coordinate space using a stereotactic neuronavigation system (BrainSight, Rogue Research Inc.).

#### MRI data analysis

MRI data was analyzed using the FieldTrip^107^, freesurfer^108^ and Connectome Workbench^109^. Individual high-resolution T1 MRI was used to construct volume conduction models via single-shell head corrected-sphere approach. Source models and surface labels from the HCP-MMP atlas^109^ were created for each participant using inner-skull and pial layer surfaces.

#### MEG data analysis

MEG data was analyzed using the FieldTrip^107^, EEGLAB^110^ and SpectralEvents (https://github.com/jonescompneurolab/SpectralEvents) toolboxes, and custom analysis scripts developed in MATLAB 2023b.

#### Pre-processing

Raw MEG data were high-pass filtered at 0.1 Hz with fourth order non-causal Butterworth filter. Powerline noise (60 Hz) and its harmonics (120 Hz) were removed by spectrum interpolation implemented in Fieldtrip. We then fitted Principal Component Analysis (PCA) using the Fieldtrip’s implementation, and Independent Component Analysis (ICA) using EEGLAB’s implementation of extended Infomax to the first 40 PCs. Spatial topographies and time series of all 40 ICs were visually inspected and those determined to be strongly associated with known electrocardiogram, eye movements and blinks artifacts were removed. MEG activity was then reconstructed using retained independent components and all other principal components, which were not used for ICA analysis.

#### Source reconstruction and parcellation

Source reconstruction of MEG data was performed using the standard pipeline in Fieldtrip. Individual volume conduction model calculated from individual high-resolution T1 MRI was used to generate the forward solution at each source dipole location. The linearly-constrained minimum-variance (LCMV) beamformer was used to compute inverse-solution from sensor-space MEG activity. The individual inverse solution data covariance matrix was obtained by averaging the mean covariance matrices of practice and rest intervals. Specifically, sensor-space MEG covariance was calculated for each 10 second practice and rest interval and averaged over trials. The individual sample noise covariance matrix was computed from 6-mins of individual pre-training rest MEG data recorded during the same session. A total of 15,684 surface-based cortical dipoles were estimated. The individual HCP-MMP atlas model, which defines 180 parcellated cortical areas per hemisphere was used to extract source-space parcellated MEG activity by averaging activity within each cortical area.

#### Theta-gamma phase amplitude coupling

We measured the strength of PAC for a wide set of amplitude frequency ranges using the modulation index (MI) as introduced previously ^111^. We first filtered (using pop_eegfiltnew.m from EEGLAB) continuous source-space parcellated MEG signal within the respective frequency bands of interest (θ=4-8Hz and γ=30-110Hz with a step of 5Hz and a window of ±6Hz; **Figure S6C**). The frequency window of γ band was chosen such that it constituted twice the center frequency of the phase signal. This procedure ensures that the side peaks that arise if the amplitude signal is modulated by a lower-frequency phase signal are included ^112^. We then extracted the instantaneous phase from the lower-frequency signal and the amplitude from the higher-frequency signal using the Hilbert transform.

During the in-lab experiment, subjects’ HIS segments started at the beginning of each practice trial and finished 4.10±0.11 s later (**Figure S5**). Thus, we extracted the time window of 0-4 s (HIS segment) and 4-10 s (post-HIS segment) relative to practice trial onset. We additionally analyzed the time window of 4-8 s to control for the time window length between HIS and post-HIS segments (**Figure S6A**). Next, we concatenated the phase and amplitude signals for each practice trial and time window (18 phase bins) and computed the modulation index (MI) following established methods ^113^. To standardize MI values across cortical areas, amplitude frequencies, and time windows, we generated 200 surrogate MIs by circularly shifting the theta-phase time series by a random interval greater than 600 samples (1s at the given sampling rate), producing a null distribution of MI values. A normal distribution was then fit to each surrogate set (*normfit*.*m*) to estimate its mean and standard deviation, which were used to z-transform the raw MI values. This normalization procedure controls for potential confounds related to amplitude power, phase properties, or evoked activity due to the transition from a rest to a practice interval that might otherwise introduce spurious differences in PAC^114^. The standardized MI values therefore permit valid comparisons across time windows and amplitude frequencies^112^. Cortical parcels exhibiting significant PAC were identified as those with normalized MI values significantly greater than zero (*t*-test, cluster-based permutation correction). Finally, θ/γ PAC values within each significant cluster were averaged for subsequent analyses.

#### Oscillatory bursting analysis

Spectral burst analysis was conducted in both sensor and source (parcellated) space MEG data, following previously established methods^115^. Transient high-power bursts were detected and characterized using the SpectralEvents Toolbox, which identifies *spectral events* as local maxima in time–frequency power exceeding a specified threshold. The analysis was event-overlap–agnostic, with the detection threshold set at six times the median power (6×factors-of-the-median, FOM) across time for each frequency bin^97^. Bursts were detected across 1-40 Hz in a 1-Hz step from continuous MEG recordings spanning the entire task period (from the start of the first to the end of the last practice trial). Each detected event was characterized by its peak time, power, central frequency, and duration (**Figure S7**). Event rate was defined as the number of detected bursts per second within HIS and post-HIS segments. Event duration was computed as the full-width at half-maximum (FWHM) of the power distribution in both the time and frequency domains, corresponding to the region where power exceeded half of the event’s local maximum. Following the recommended procedure used in the previous studies^115^ the first 1 s of each practice interval was excluded from analysis to minimize contamination from transient spectral activity associated with practice onset.

## QUANTIFICATION AND STATISTICAL ANALYSIS

We assessed the significance of several variables during the HIS segment, including the number of keypresses, keypress rate, number of chunks, chunk rate, chunk content and HIS duration. We compared these metrics across different groups and as practice progressed within each skill group. To perform the across-group comparison, we used a one-way ANOVA (*anova1*.*m*) followed by Tukey’s honestly significant difference procedure for post-hoc analysis (*multcompare*.*m*). The progression of these values with practice trials was evaluated using linear-mixed effects models (LMM, *fitlme*.*m*). The choice of an LMM was made to accommodate inter-individual differences by including them as a random effect. All models were fitted using restricted maximum likelihood estimation. We tested the significance of fixed effects in the model using a Type III analysis of variance (ANOVA) with Satterthwaite’s method (*fixedEffects*.*m* and *anova*.*m*). For behavioral evaluations, we used trial number as the dependent variable to measure trial-by-trial development of variables in HIS segments. On the other hand, within-cluster averaged θ/γ PAC was the dependent variable when predicting the number of HIS contents.

During the MEG analysis, we used a cluster-based multiple comparison correction approach to identify frequencies of interest, spatial contribution of parcellated brain regions to θ/γ PAC and β bursts^116^. This correction approach was chosen to address the multiple comparisons problem inherent in the high spatial resolution of the HCP-MMP atlas, while effectively leveraging its specific parcellation of 360 cortical brain regions.

## KEY RESOURCE TABLE

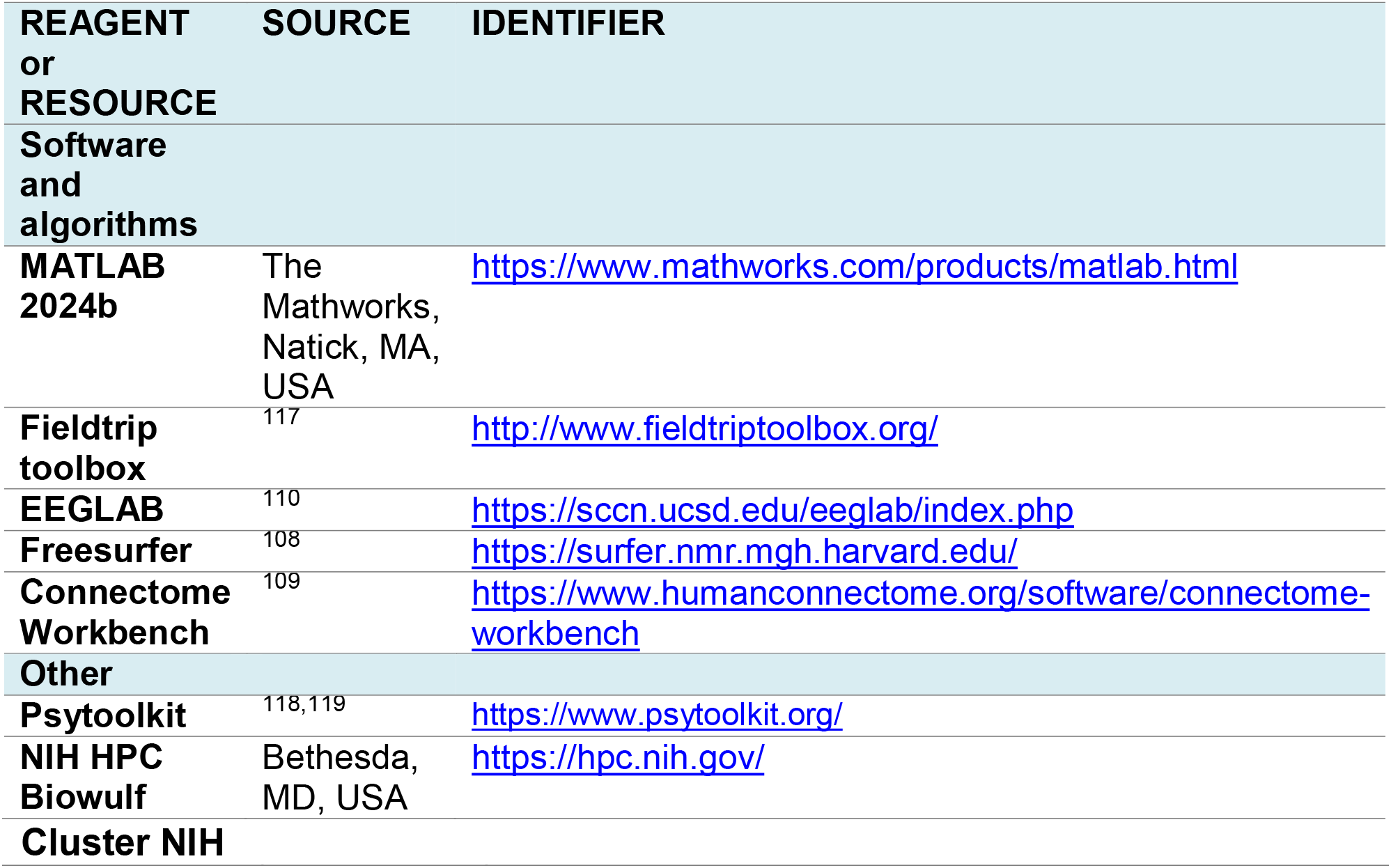

## Acknowledgments

We thank Ms. Catherine Blumhorst and NIMH MEG Core Facility staff for their support. This work utilized the computational resources of the NIH HPC Biowulf cluster (http://hpc.nih.gov). This research was supported by the Intramural Research Program of the National Institutes of Health (NIH). FI was partly supported by KAITOKU fellowship funded by the Japan Society for the Promotion of Science (JSPS). The contributions of the NIH author(s) are considered Works of the United States Government. The findings and conclusions presented in this paper are those of the author(s) and do not necessarily reflect the views of the NIH or the U.S. Department of Health and Human Services.

## Author Contributions

Conceptualization: F.I., E.R.B. and L.G.C.; Methodology: F.I., E.R.B. and L.G.C.; Formal Analysis: F.I., E.R.B. and L.G.C.; Investigation: F.I., E.R.B. and L.G.C.; Writing—original draft: F.I., E.R.B. and L.G.C.; Writing—review & editing: F.I., W.H., D.K., E.R.B. and L.G.C.; Visualization: F.I., E.R.B. and L.G.C.; Supervision: E.R.B. and L.G.C.

## Declaration of generative AI and AI-assisted technologies

During the preparation of this work the authors used GPT-5.2 for editing purposes. After using this tool/service, the authors reviewed and edited the content as needed and take full responsibility for the content of the published article.

## Supplementary data

**Figure S1.**
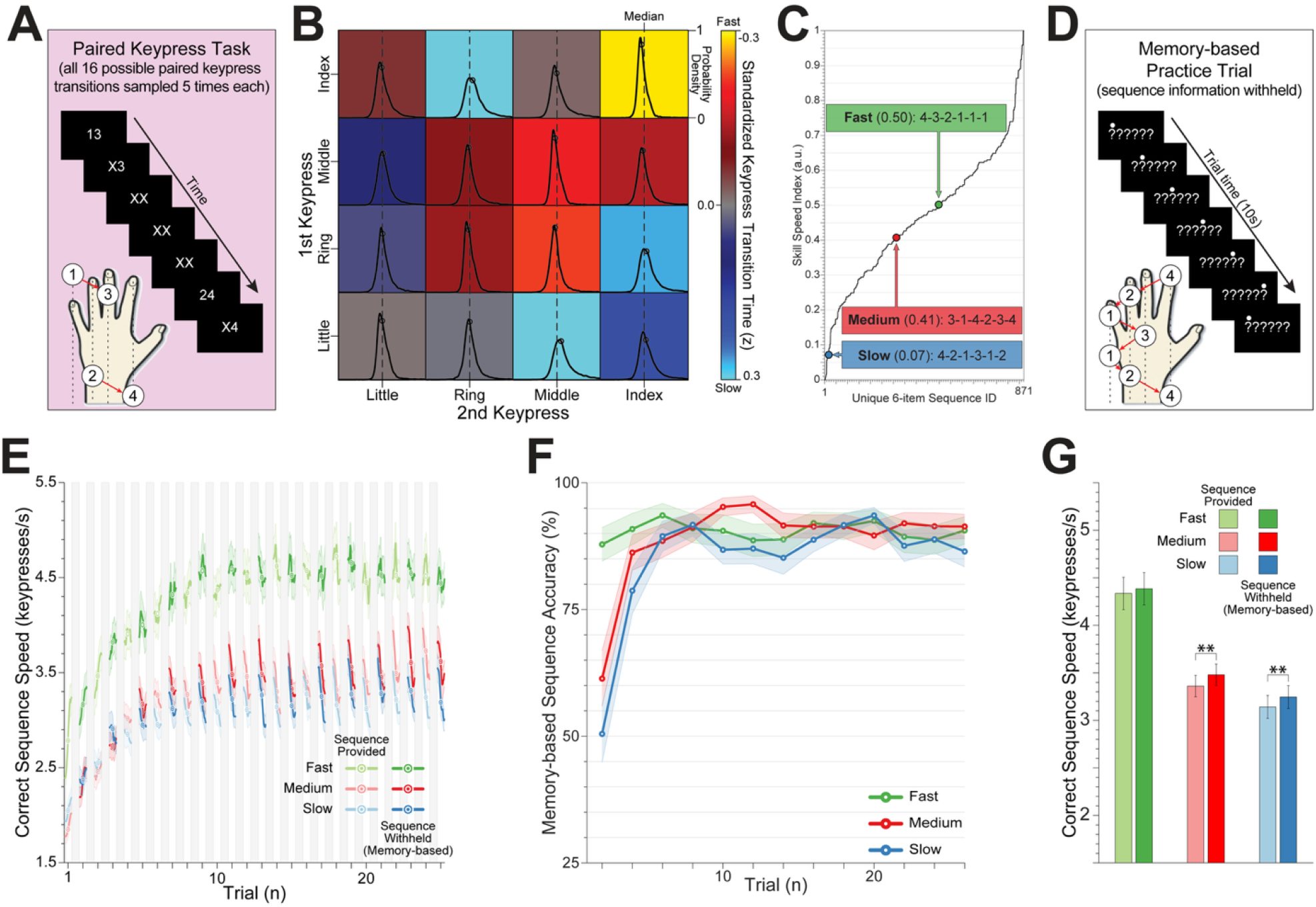
Rapid emergence of internal sequence representations during early practice related to Figure 1. (**A**) We first collected a large independent dataset of keypress transition times (KTTs) from healthy volunteers (N=2152) performing all 16 possible two-finger transitions using the little, ring, middle, and index fingers of their non-dominant (left) hand on a standard keyboard. On each trial, participants saw a pair of numeric cues (e.g., “24”) indicating the required finger pair (1 = little, 2 = ring, 3 = middle, 4 = index) and were instructed to respond as quickly and accurately as possible. Individual numbers were replaced with “X” as the corresponding keypresses were executed. Incorrect responses triggered a repetition of the trial until a correct response was registered. The inter-trial interval was uniformly jittered between 200–500 ms, and each of the 16 possible transitions were sampled 5 times in pseudorandom order. (**B**) For each participant, transition times separating keypress pairs (KTT_pairs_) were z-normalized, and a group-level matrix of median z-score shifts for each keypress pair was constructed. (**C**) We then used this matrix to synthesize predictive estimates of the relative duration of all possible 6-item keypress sequences by summing their constituent KTT_pairs_, thereby deriving a Skill Speed Index (SSI) that characterizes the predicted execution speed of every possible 6-item keypress sequence. Three skills of different speeds are shown. **(D-G)**. In a separate experiment, memorability was assessed for each of the three sequence skills. Participants in each speed group completed 26 practice trials with alternating task demands: on odd-numbered trials, the sequence remained visible to allow visually guided practice; on even-numbered trials, the cues were removed and participants reproduced the sequence from memory **(D)**. Memory-guided performance therefore served as an index of the emergence of internal sequence representations. **(E)** Across all three skills, internal representations emerged rapidly during early practice, with most participants able to produce the sequence from memory by trial 6. **(F)** Proportion of sequences correctly typed from memory on even-numbered trials. For the fast skill, participants reached >95% of asymptotic memory-based accuracy by trial 2, whereas the slower, more difficult skills required 4 and 6 trials, respectively, to reach comparable levels. These differences in memorability may have contributed to across-skill variation in chunk structure, including the greater number of keypresses per chunk for the fast relative to the slow skill (**Figure 2F**). (**G**) Once the sequence had been memorized, mean execution speed was higher on trials performed without visual stimuli (internally generated) than on those with visual stimuli (BHFDR-corrected paired t-tests: Fast: t_61_=1.10, p=0.27; Medium: t_68_=3.54, p=0.02; Slow: t_69_=3.10, p=0.04).

**Figure S2.**
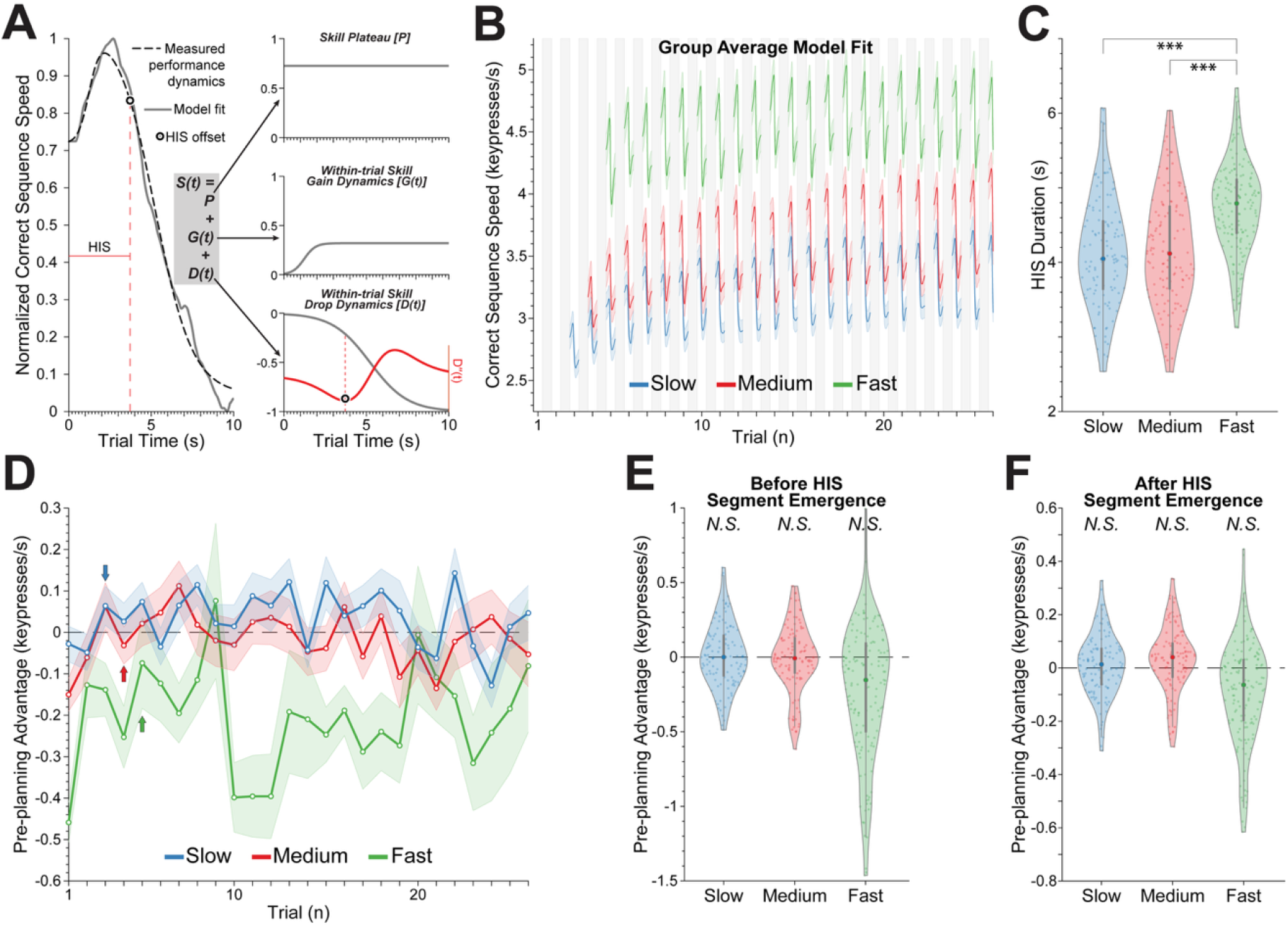
Modelling within-trial skill dynamics related to Figures 1 and 2. (**A**) Skill dynamics (*S(t);* shown for representative individual subject trial) over each practice trial were modeled by fitting a double-sigmoidal function^41^ that included skill plateau (*P*), within-trial gain (*G(t)*) and within-trial drop (*D(t)*) dynamics. The end of the HIS segment was defined by the maximum deceleration point (minimum of the 2^nd^ derivative) of within-trial drop dynamics (see **Equation 3** in Methods). (**B**) Within each skill group, subjects’ individual trial models were aligned on the HIS segment offset and averaged (as shown for the last two trials in **Fig 1C-D**). Note the stability of this pattern following the emergence of HIS segments (trials 3, 4 and 5 for slow, medium and fast skills respectively). **(C)** The duration of the HIS segment was longer for the fastest skill than the slower skills. The duration of HIS segments were stable throughout the practice (Slow: F_1,2750_=0.28, *p*=0.99; Medium: F_1,2787_=2.57×10^-4^, p=0.99; Fast: F_1,3357_=1.75×10^-3^, p=0.99), resulting in training dependent increases in keypress rates (**Fig 2C**). **(D–F)** Pre-planning advantage in the three speed groups (Slow, Medium, and Fast), computed across the full training period **(D)**, and separately before **(E)** and after **(F)** the emergence of HIS segments. In **(D)**, arrows mark the time-point at which HIS segments first emerged in each speed group. Negative values indicate faster execution in the second sequence than in the first, whereas positive values indicate faster execution in the first sequence, indicative of a pre-planning advantage. **(E)** Before HIS emergence, participants showed speeding from the first to the second sequence across all groups, arguing against a pre-planning boost (BHFDR-corrected t-test against zero; Slow: t_94_=-1.07, p=1.0; Medium: t_116_=-0.55, p=1.0; Fast: t_153_=-5.77, p=1.00). **(F)** After HIS emergence, speed differences were around zero, indicating either continued speeding in the second sequence or no difference between sequences, again inconsistent with a pre-planning advantage (Slow: t_121_=1.56, p=0.18; Medium: t_126_=-0.40, p=0. 79; Fast: t_156_=-1.96, p=0.97).

**Figure S3.**
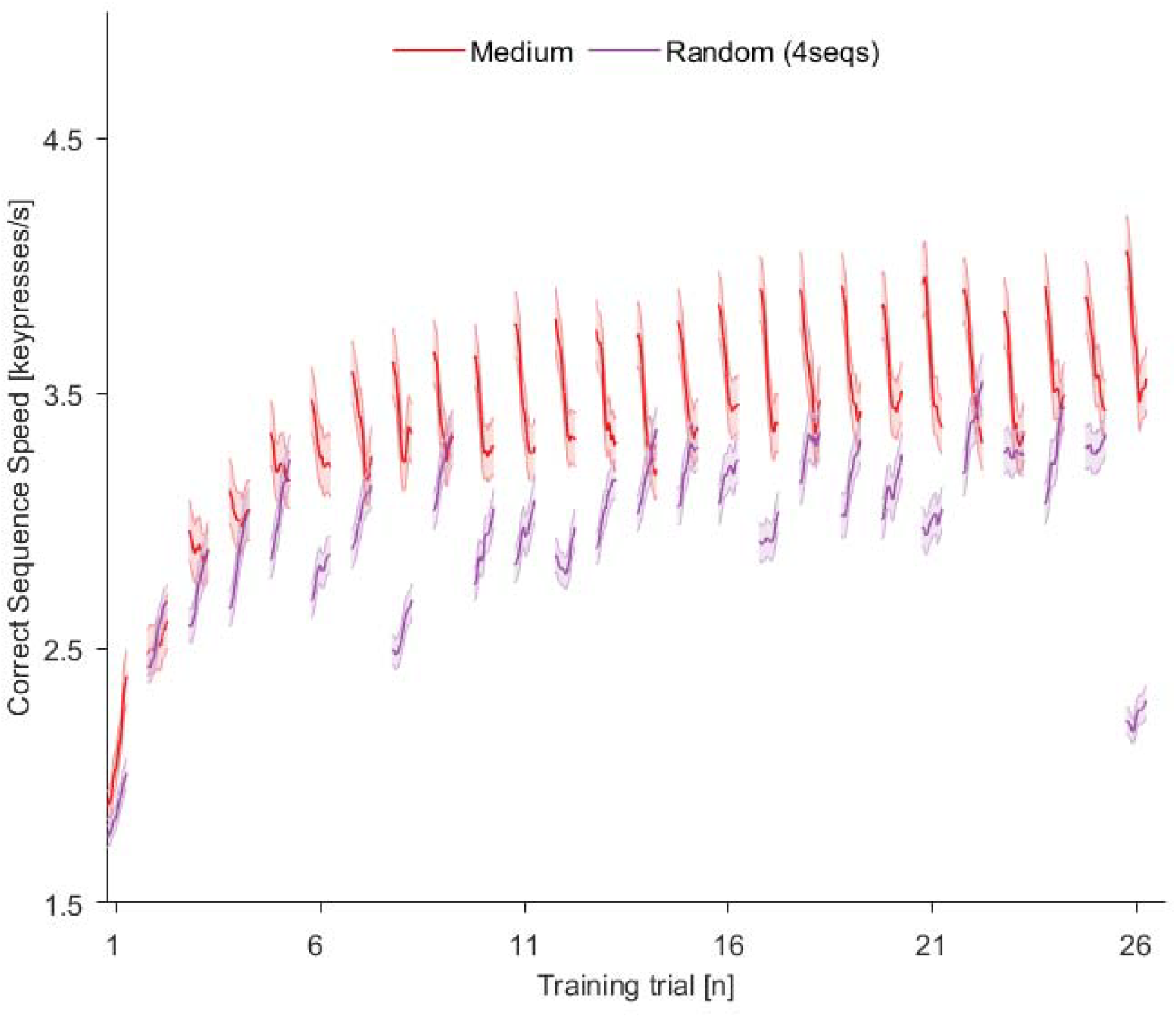
Learning dependence of HIS segments related to Figure 1. An additional cohort (N=156) practiced four non-repeating sequences in pseudorandom order from trials 2–25, with the same medium-speed sequence performed on trials 1 and 26. Despite comparable execution speed, intervening practice on non-repeating sequences (a) produced negligible learning of the medium sequence (first vs. last purple trials), in contrast to robust gains with repeated-sequence practice (red trials), and (b) failed to generate HIS segments. Thus, non-repetitive practice at similar speeds yielded qualitatively different within-trial dynamics, arguing against speed-related fatigue as the origin of HIS segment structure.

**Figure S4.**
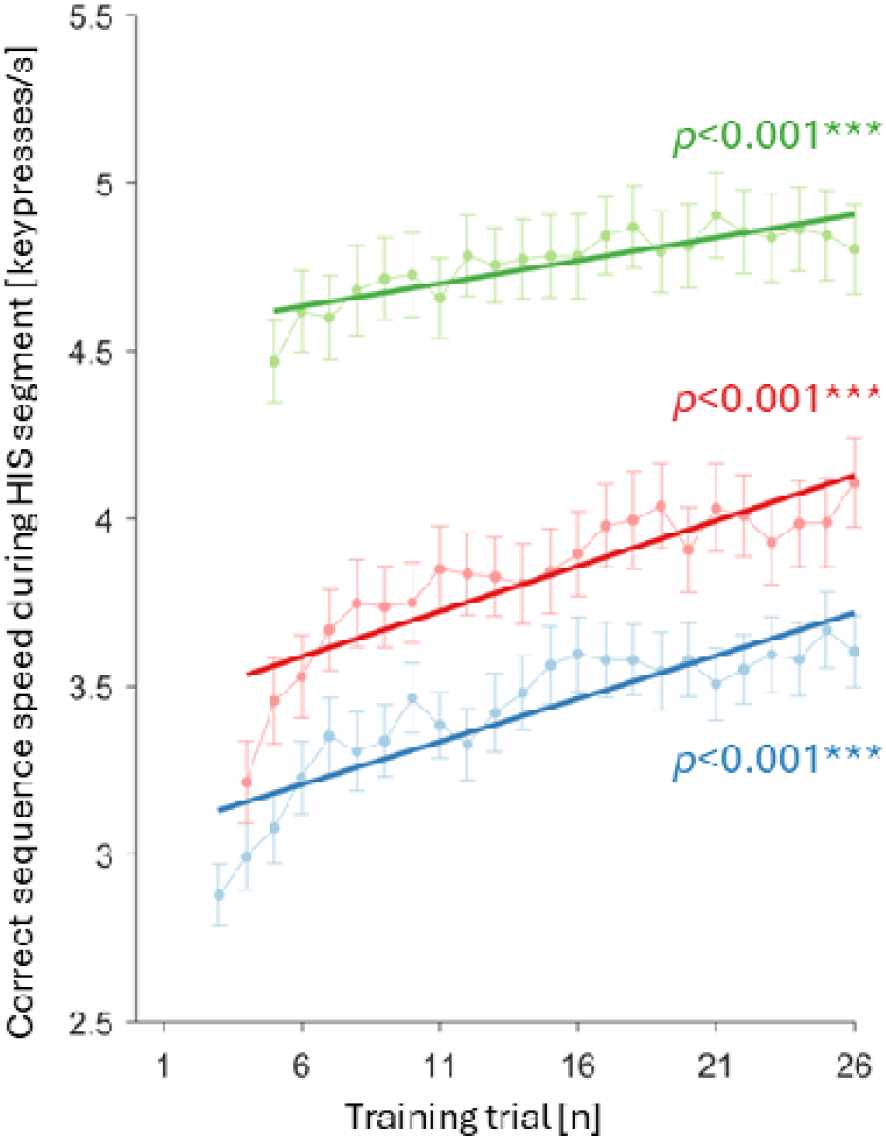
Skill performance during HIS segments related to Figure 2. Skill speed during HIS segments increased progressively with practice in all groups after HIS emergence and the increase in speed persisted even after median trial performance speed reached plateau (trials 6-26; BHFDR-corrected LMMs; Slow, F_1,2290_=70.98, p<0.001; Medium, F_1,2429_=116.34, p<0.001; Fast, F_1,3044_=29.82, p<0.001).

**Figure S5.**
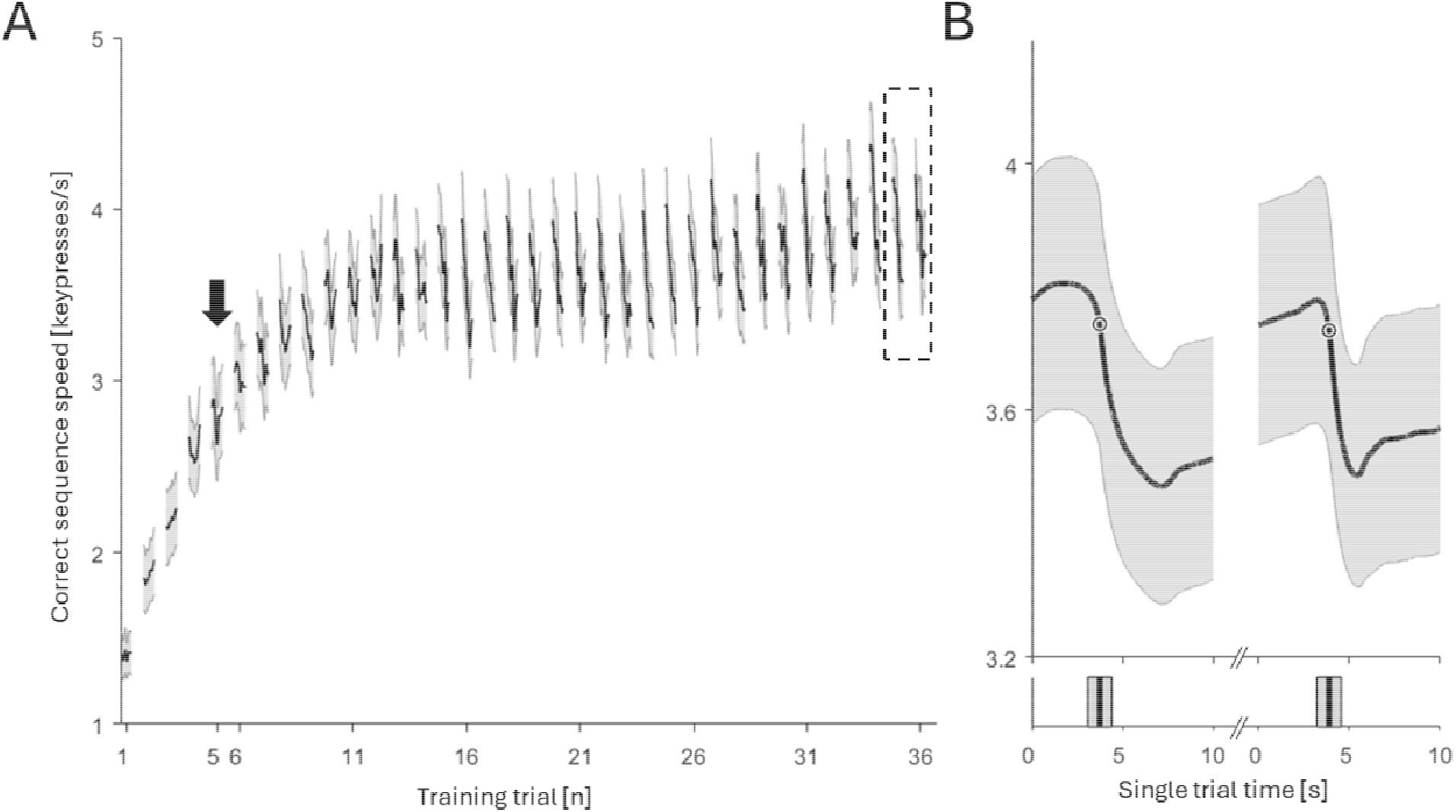
Replication of HIS dynamics in an independent MEG dataset related to Figure 1. In Experiment 2, we examined MEG oscillatory dynamics in participants learning a medium-speed sequential skill over 36 practice trials (N = 32; https://openneuro.org/datasets/ds006502/versions/1.0.0). **(A)** Performance curves, computed as in Experiment 1, revealed the emergence of high-initial-skill (HIS) segment dynamics around trial 5 (arrow), closely paralleling the onset observed previously (arrows in **Fig 1B**). **(B)** Group-level intra-trial performance profiles for the last two trials (35 and 36), aligned to the end of the HIS segment and averaged as in Experiment 1, show the same characteristic within-trial pattern. Thus, the HIS segments intra-trial dynamics described in Experiment 1 were reproduced in an independent cohort and recording environment.

**Figure S6.**
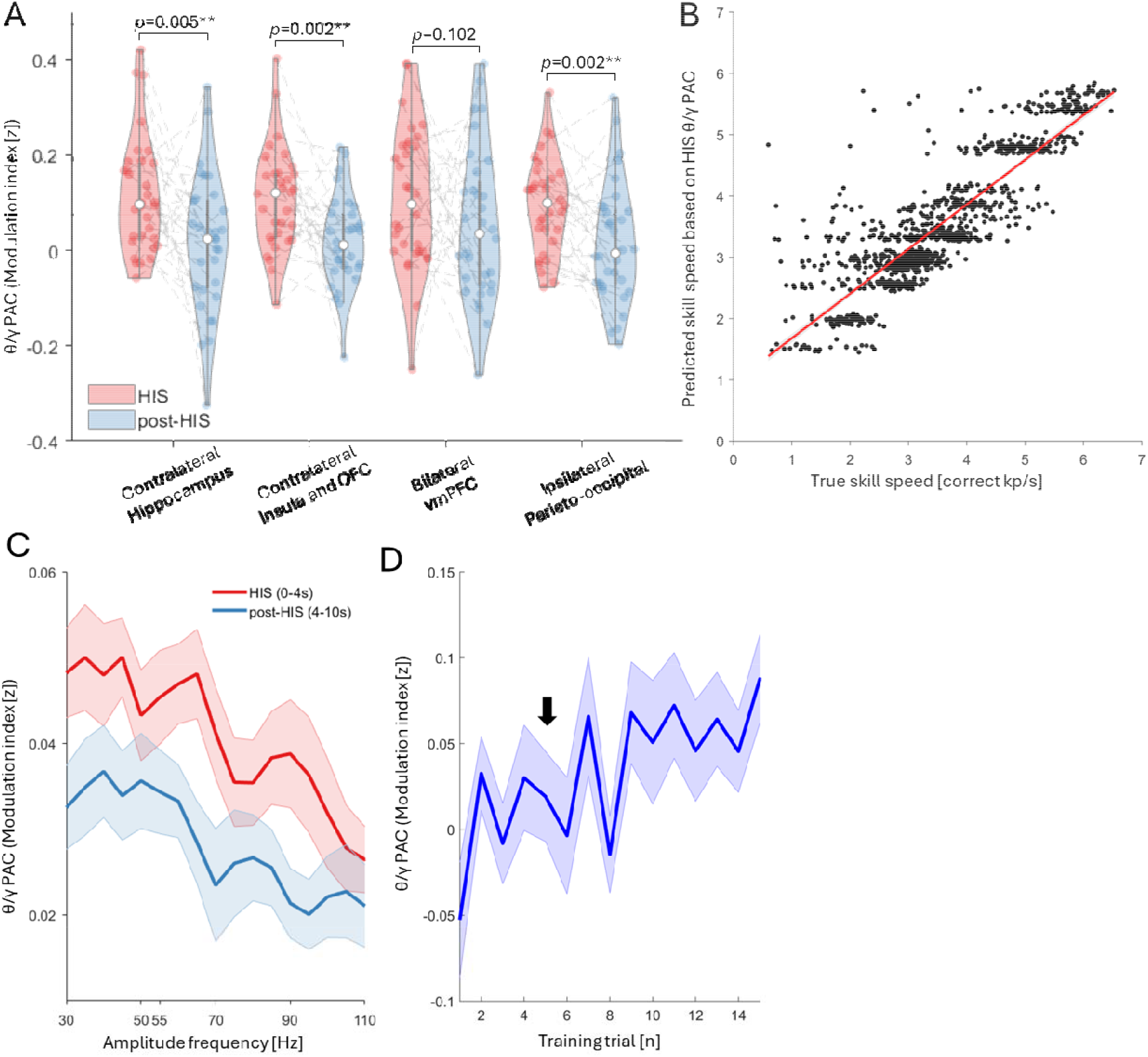
Complementary θ/γ PAC analyses related to Figure 3. **(A)** We compared θ/γ PAC during HIS segments (0–4 s) with duration-matched θ/γ PAC during the immediately following interval (4–8 s). The resulting spatial pattern closely matched that obtained in **Fig 3C**, which used a slightly longer post-HIS window (4–10 s). No significant predictive effects were observed in the other regions (contralateral insula/OFC: *t*_986_==1.29, *p*=0.39; bilateral vmPFC: *t*_985_=−0.36, *p*=0.72; ipsilateral parieto-occipital: *t*_987_=0.44, *p*=0.72). (**B**) Hippocampal θ/γ PAC during HIS segments was used to model skill performance (mean correct sequence speed) using linear mixed-effects models. The modeled speeds derived from hippocampal θ/γ PAC (y-axis) closely tracked the observed mean skill speeds of practice trials (x-axis), indicating that hippocampal θ/γ PAC reliably predicted skill level across practice. (**C**) Whole-brain source-space θ/γ PAC strength, parcellated across regions and normalized by 200 time-shifted surrogates, was quantified during HIS and post-HIS segments (mean ± SEM; θ = 4–8 Hz, γ = 30–110 Hz in 5-Hz steps with ±6-Hz windows; trials 5–36). Because θ/γ PAC was strongest in the 30–55 Hz range, this γ-band interval was used for all analyses. (**D**) Whole-brain source-space θ/γ PAC within the time window encompassing HIS segments (0-4 s) increased progressively with practice (trial 1-15; LMM, F_1,478_=12.8, *p*<0.001).

**Figure S7.**
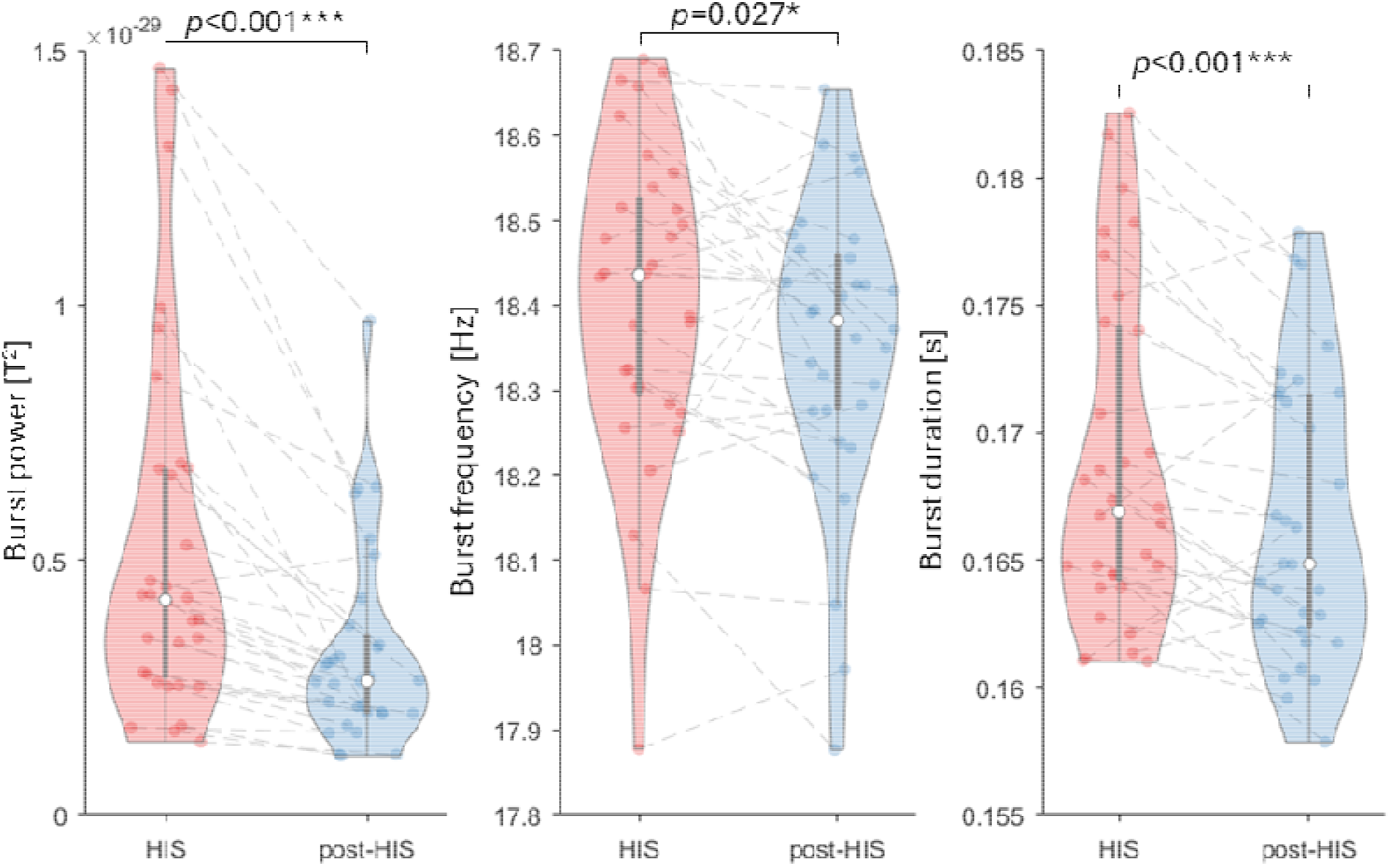
related to Figure 4: β band burst activity during HIS segments exhibited higher power, higher frequency, and longer duration than during post-HIS segments (paired t-tests). HIS segments were determined in each practice trial and each participant.

## Supplementary Tables

**Table S1:** Our analysis identified four clustered brain regions contributing to θ/γ PAC difference during HIS segments relative to its time-shifted surrogates (n=200). Tables below document the subregions defined in HCP-MMP atlas contained within each cluster.

(A) Contralateral hippocampal cortex
| Hemisphere<br>("L" or "R") | Area Name | Area Description | $t_{31}$ value | $p$ value |
| --- | --- | --- | --- | --- |
| R | H | Hippocampus | 2.76 | $3.76 \times 10^{-3}$ |
| R | PHA1 | ParaHippocampal area 1 | 3.58 | $5.77 \times 10^{-4}$ |
| R | PHA2 | ParaHippocampal area 2 | 3.98 | $1.91 \times 10^{-4}$ |
| R | PHA3 | ParaHippocampal area 3 | 4.07 | $1.52 \times 10^{-4}$ |

(B) Contralateral insula and orbitofrontal cortex
| Hemisphere<br>("L" or "R") | Area Name | Area Description | $t_{31}$ value | $p$ value |
| --- | --- | --- | --- | --- |
| R | 13l | Area 13l | 3.27 | $1.30 \times 10^{-3}$ |
| R | OFC | Orbitofrontal cortex | 3.92 | $2.29 \times 10^{-4}$ |
| R | Pir | Piriform cortex | 2.79 | $4.46 \times 10^{-3}$ |
| R | AVI | Anterior Ventral Insular Area | 2.87 | $3.71 \times 10^{-3}$ |
| R | AAIC | Anterior Agranular Insula Complex | 3.39 | $9.56 \times 10^{-4}$ |
| R | TGd | Area TG dorsal | 3.39 | $9.58 \times 10^{-4}$ |

(C) Bilateral vmPFC
| Hemisphere<br>("L" or "R") | Area Name | Area Description | $t_{31}$ value | $p$ value |
| --- | --- | --- | --- | --- |
| L | p32 | Area p32 prime | 2.88 | $3.58 \times 10^{-3}$ |
| L | 10r | Area 10r | 3.17 | $1.69 \times 10^{-3}$ |
| R | 10r | Area 10r | 3.30 | $1.22 \times 10^{-3}$ |

**(D)** Ipsilateral parieto-occipital regions
| Hemisphere<br>("L" or "R") | Area Name | Area Description | $t_{31}$ value | $p$ value |
| --- | --- | --- | --- | --- |
| L | V3A | Area V3A | 3.49 | $7.30 \times 10^{-4}$ |
| L | POS2 | Parieto-Occipital Sulcus Area 2 | 2.88 | $3.53 \times 10^{-3}$ |
| L | IPS1 | IntraParietal Sulcus Area 1 | 3.03 | $2.45 \times 10^{-3}$ |
| L | DVT | Dorsal Transitional Visual Area | 2.93 | $3.12 \times 10^{-3}$ |
| L | V6A | Area V6A | 2.91 | $3.31 \times 10^{-3}$ |

**Table S2:** Our analysis identified three clustered brain regions contributing to β burst rate difference between HIS and post-HIS segments. Tables below document the subregions defined in HCP-MMP atlas contained within each cluster.

| Hemisphere<br>("L" or "R") | Area Name | Area Description | $t_{31}$ value | $p$ value |
| --- | --- | --- | --- | --- |
| R | PEF | Premotor Eye Fields | 3.25 | $1.39 \times 10^{-3}$ |
| R | 6v | Ventral Area 6 | 3.64 | $4.93 \times 10^{-4}$ |
| R | 8C | Area 8C | 2.77 | $4.73 \times 10^{-3}$ |
| R | 44 | Area 44 | 2.72 | $5.24 \times 10^{-3}$ |
| R | 45 | Area 45 | 3.50 | $5.25 \times 10^{-4}$ |
| R | 47l | Area 47l | 2.57 | $7.58 \times 10^{-3}$ |
| R | 6r | Rostral Area 6 | 2.94 | $3.04 \times 10^{-3}$ |
| R | IFJa | Area IFJa | 2.89 | $3.48 \times 10^{-3}$ |
| R | IFJp | Area IFJp | 2.94 | $3.11 \times 10^{-3}$ |
| R | IFSp | Area IFSp | 4.21 | $1.02 \times 10^{-4}$ |
| R | IFSa | Area IFSa | 3.59 | $5.63 \times 10^{-4}$ |
| R | p9-46v | Area posterior 9-46v | 2.85 | $3.86 \times 10^{-3}$ |
| R | 47s | Area 47s | 2.55 | $7.98 \times 10^{-3}$ |
| R | Pol2 | Posterior Insula area 2 | 3.04 | $2.42 \times 10^{-3}$ |
| R | TA2 | Area TA2 | 3.52 | $6.78 \times 10^{-4}$ |
| R | FOP4 | Frontal Opercular Area 4 | 3.76 | $3.53 \times 10^{-4}$ |
| R | MI | Middle Insular Area | 3.04 | $2.36 \times 10^{-3}$ |
| R | Pir | Piriform cortex | 3.07 | $2.21 \times 10^{-3}$ |
| R | AVI | Anterior Ventral Insular Area | 2.58 | $7.51 \times 10^{-3}$ |
| R | AAIC | Anterior Agranular Insula Complex | 4.42 | $5.57 \times 10^{-5}$ |
| R | FOP1 | Frontal Opercular Area 1 | 3.53 | $6.65 \times 10^{-4}$ |
| R | FOP3 | Frontal Opercular Area 3 | 3.53 | $6.63 \times 10^{-4}$ |
| R | STGa | Area STGa | 5.81 | $1.07 \times 10^{-6}$ |
| R | A5 | Auditory 5 Complex | 3.25 | $1.38 \times 10^{-3}$ |
| R | STSda | Area STSd anterior | 4.25 | $9.06 \times 10^{-5}$ |
| R | TGd | Area TG dorsal | 3.33 | $1.13 \times 10^{-3}$ |
| R | TE2a | Area TE2 anterior | 3,86 | $2.68 \times 10^{-4}$ |
| R | Pol1 | Area Posterior Insular 1 | 3.11 | $1.99 \times 10^{-3}$ |
| R | FOP5 | Frontal Opercular Area 5 | 3.51 | $7.00 \times 10^{-4}$ |
| R | p47r | Area posterior 47r | 3.76 | $3.55 \times 10^{-4}$ |
| R | STSva | Area STSv anterior | 2.92 | $3.18 \times 10^{-3}$ |
| R | TE1m | Area TE1 Middle | 2.54 | $8.13 \times 10^{-3}$ |
| R | PI | Para-Insular Area | 3.70 | $4.23 \times 10^{-4}$ |

(B) Bilateral anterior cingulate cortex
| Hemisphere<br>("L" or "R") | Area Name | Area Description | $t_{31}$ value | $p$ value |
| --- | --- | --- | --- | --- |
| L | d32 | Area dorsal 32 | 3.10 | $2.07 \times 10^{-3}$ |
| L | 8BM | Area 8BM | 3.69 | $4.26 \times 10^{-4}$ |
| R | a24pr | Anterior 24 prime | 3.70 | $4.09 \times 10^{-4}$ |
| R | p32pr | Area p32 prime | 2.47 | $9.51 \times 10^{-3}$ |
| R | d32 | Area dorsal 32 | 3.59 | $5.60 \times 10^{-4}$ |
| R | 8BM | Area 8BM | 2.80 | $4.41 \times 10^{-3}$ |
| R | a32pr | Area anterior 32 prime | 3.27 | $1.33 \times 10^{-3}$ |
| R | p24 | Area posterior 24 | 3.04 | $2.41 \times 10^{-3}$ |

(C) Ipsilateral insula, frontal and temporal cortex
| Hemisphere<br>("L" or "R") | Area Name | Area Description | $t_{31}$ value | $p$ value |
| --- | --- | --- | --- | --- |
| L | A1 | Primary Auditory Cortex | 4.37 | $6.47 \times 10^{-5}$ |
| L | 44 | Area 44 | 2.59 | $7.17 \times 10^{-3}$ |
| L | 43 | Area 43 | 2.71 | $5.41 \times 10^{-3}$ |
| L | OP4 | Area OP4/PV | 3.44 | $8.41 \times 10^{-4}$ |
| L | OP1 | Area OP1/SII | 2.95 | $3.01 \times 10^{-3}$ |
| L | OP2-3 | Area OP2-3/VS | 3.90 | $2.41 \times 10^{-4}$ |
| L | 52 | Area 52 | 4.41 | $5.72 \times 10^{-5}$ |
| L | Pol2 | Posterior Insular Area 2 | 4.04 | $1.66 \times 10^{-4}$ |
| L | TA2 | Area TA2 | 3.54 | $6.44 \times 10^{-4}$ |
| L | FOP1 | Frontal Opercular Area 1 | 3.43 | $8.50 \times 10^{-4}$ |
| L | FOP2 | Frontal Opercular Area 2 | 3.14 | $1.86 \times 10^{-3}$ |
| L | STGa | Area STGa | 4.42 | $5.57 \times 10^{-5}$ |
| L | PBelt | ParaBelt Complex | 2.51 | $8.65 \times 10^{-3}$ |
| L | A5 | Auditory 5 Complex | 3.40 | $9.36 \times 10^{-4}$ |
| L | STSda | Area STSd anterior | 4.83 | $1.73 \times 10^{-5}$ |
| L | TE1a | Area TE1 anterior | 2.57 | $7.62 \times 10^{-3}$ |
| L | PFop | Area PF opercular | 4.29 | $8.06 \times 10^{-5}$ |
| L | Pol1 | Area Posterior Insular 1 | 3.12 | $1.96 \times 10^{-3}$ |
| L | Ig | Insular Granular Complex | 3.24 | $1.41 \times 10^{-3}$ |
| L | MBelt | Medial Belt Complex | 4.04 | $1.64 \times 10^{-4}$ |
| L | A4 | Auditory 4 Complex | 3.00 | $2.65 \times 10^{-3}$ |
| L | STSva | Area STSv anterior | 4.15 | $1.19 \times 10^{-4}$ |
| L | TE1m | Area TE1 Middle | 2.51 | $8.66 \times 10^{-3}$ |
| L | PI | Para-Insular Area | 2.46 | $9.75 \times 10^{-3}$ |

